# Lemur gut microeukaryotic community variation is not associated with host phylogeny, diet, or habitat

**DOI:** 10.1101/2023.01.17.524408

**Authors:** Mariah E. Donohue, Zoe L. Hert, Carly E. Karrick, Amanda K. Rowe, Patricia C. Wright, Lovasoa J. Randriamanandaza, François Zakamanana, Stela Nomenjanahary, Kathryn M. Everson, David W. Weisrock

## Abstract

Gut prokaryotic (GP) community variation is often associated with host evolutionary and ecological variables; whether these factors drive variation in other gut taxa remains largely untested. We present a one-to-one comparison of GP (16S rRNA metabarcoding) and microeukaryotic (GME) (18S rRNA metabarcoding) community patterning among 12 species of lemurs. Lemurs were sampled from dry forests and rainforests of southeastern Madagascar and display a range of phylogenetic and ecological diversity. We found that while lemur GPs vary with host taxonomy, diet, and habitat, GMEs have no association with these factors. As a mechanism, we suggest purifying selection purges microbes with negative and commensal associations, while positive selection promotes the persistence of beneficial microbes. It is therefore likely that a greater proportion of GMEs comprise taxa with commensal, transient, and parasitic symbioses compared with GPs, many of which are mutualists. Our study reveals different microbial taxa are shaped by unique selective pressures.

## INTRODUCTION

Mammalian gut microbiomes comprise diverse communities of bacteria, archaea, fungi, worms, protists, and viruses, which perform specialized functions critical to the health, ecology, and evolution of their hosts. At the organismal level, gut microbes can both cause and fight infections (Rosa et al., 2018), improve nutrient acquisition in response to changing diets (David et al., 2014), and promote healthy neurological functioning (Montiel-Castro et al., 2013). At the host population and species levels, they are known to influence major evolutionary processes, including ecological adaptation (Alberdi et al., 2016; Greene et al., 2020; Wagener et al., 2020) and speciation (at least in insects; Sharon et al., 2011; Brucker & Bordenstein, 2013; Rennison et al., 2019; Miller et al., 2021). Gut microbiomes also demonstrate intra- and inter-species variation associated with host ecological and evolutionary forces. However, most research in this field has solely described variation in gut prokaryotic communities (GPs; bacteria and archaea), creating a major gap in our understanding of variation among communities of gut microeukaryotes (GMEs; fungi, worms, and protists) and inter-kingdom interactions (i.e., interactions between GPs and GMEs).

GMEs help regulate gut microbiome function through a variety of ecological roles, including decomposition, primary production, predation, and parasitism (Keeling & del Campo, 2017). Despite such a diversity of ecological niches, researchers have primarily studied GMEs through the lens of infectious disease and parasitology. This bias stems from the fact that GMEs account for an overwhelming majority of disease-causing parasites, including *Plasmodium falciparum, Toxoplasma gondii, Giardia duodenalis*, and *Entamoeba histolytica* (Walker et al., 2011). Similarly, before the advent of next-generation metabarcoding, GPs were regarded as pathogens, not hyper-diverse communities with negative, positive, and neutral host associations (Hooper & Gordon, 2001). GMEs have yet to undergo the metabarcoding renaissance that elevated our understanding of GPs from pathogens to complex communities. The most recent meta-analysis found just 5% of published microbiome studies examined GMEs (Keeling & del Campo, 2017). As a result, our present understanding of the gut microbiome overlooks the full scope of GME diversity and composition, leaving basic questions about community variation largely unexplored.

One outstanding question is whether GMEs are shaped by the same selective pressures as GPs. Host phylogeny, diet, and habitat are three major factors known to drive GP variation across the Tree of Life (e.g., Groussin et al., 2017), though their influences on GME variation are poorly understood. Wild primates represent a compelling system for initial comparisons of GP and GME communities, given their incredible ecological and taxonomic diversity and close relatedness to humans. Still, of the albeit limited studies explicitly examining primate GME community variation, a consensus about the driving forces has not been reached. For example, in a comparison of chimpanzee gut microbiome variation across central Africa, de Mesquita et al., (2021) found that GP and GME communities largely mirrored one another in mechanistic drivers, as both were significantly shaped by host genetics, diet, and ecology. Longitudinal studies of the lemurs *Eulemur rufifrons* and *Microcebus rufus* showed GME communities vary over time, likely in tandem with environmental variation (e.g., seasonality) and life history (Aivelo et al., 2015; Murillo et al., 2022). In the only multi-species comparative dataset to-date, Mann et al. (2020) sampled representative species from across the Primate order and found weak significant effects of phylogeny and diet. This finding contrasts with those focused on GPs, which tend to report evidence of strong phylosymbiosis (phylogenetic signal of the gut microbiome) (e.g., Amato et al., 2019). Because sampling scale is known to impact detection of phylosymbiosis and ecological effects in GPs (e.g., Greene et al., 2019), it is possible that patterns strengthen when analyses are narrowed to only include closely-related species with significant ecological diversity.

This study compares patterns of GP and GME community variation across multiple species of lemurs, a speciose monophyletic Primate clade endemic to the island of Madagascar, where they occupy a wide range of habitats and exhibit remarkable adaptations to harsh environments (Wright, 1999). They also demonstrate incredible ecological niche diversity, allowing for coexistence between both closely- and distantly-related species. At the same time, congeners often occupy different environmental extremes (Herrera, 2017). This dynamic ecological and evolutionary matrix sets the stage for a natural experiment to identify the major drivers of gut microbiome variation. Most endosymbiotic lemur research has focused on microscopic identification of parasites (e.g., Raharivololona & Ganzhorn, 2010; Junge et al., 2011; Lazdane et al., 2014; Rakotoniaina et al., 2016) or characterizing GP diversity (e.g., Umanets et al., 2018; Donohue et al., 2019; Greene et al., 2019; Perofsky et al., 2019; Donohue et al., 2022). Aivelo et al. (2018) used metabarcoding to describe GMEs of sympatric lemur species but did not analyze microbial patterns on the community level.

Here, we present a one-to-one comparison of lemur GP and GME composition and diversity. Our primary goal was to elucidate the causes and consequences of lemur GME variation in nature. Specifically, we addressed the following questions: (1) which GME species reside in the lemur gut? (2) which evolutionary and ecological variables drive GME community variation? and (3) how do cross-kingdom interactions (i.e., microbe-microbe interactions between GMEs and GPs) shape the lemur gut microbiome? Our study offers new, surprising insight into the factors (or lack thereof) influencing GME community patterning.

## MATERIALS AND METHODS

### Study Sites and Sample Collection

We used 72 fecal samples originally used in Donohue et al. (2022), which used 16S rRNA sequence data to test GP phylosymbiosis, and Rowe et al. (2021), which used metabarcoding to detect insectivory. Here, we used these samples to generate 18S rRNA sequence data for GME analysis. This dataset comprised 12 lemur species, spanning nine genera and three families. These species show striking variation in diet, activity patterns, body size, and sociality. Fecal samples were collected from three National Parks (Ranomafana, rainforest; Isalo, dry forest; Zombitse-Vohibasia, dry forest) in southern Madagascar (Fig. 1; Table 1). Precise sampling methodologies were described in Rowe et al. (2021) and Donohue et al. (2022).

**Table 1:** Taxonomic identification and ecological classifications of study subjects.

| Host family | Host species | # Samples | Primary foods | Habitat | Site |
| --- | --- | --- | --- | --- | --- |
| Cheirogaleidae | *Cheirogaleus medius* | 3 | Insects, fruits | Dry forest | Zombitse |
|  | *Microcebus rufus* | 15 | Insects, fruits | Rainforest | Ranomafana |
|  | *Mirza coquereli* | 5 | Insects, fruits | Dry forest | Zombitse |
|  | *Phaner pallescens* | 4 | Gum | Dry forest | Zombitse |
| Lemuridae | *Eulemur rubriventer* | 5 | Fruits | Rainforest | Ranomafana |
|  | *Eulemur rufifrons* (east) | 11 | Leaves, fruits, flowers | Rainforest | Ranomafana |
|  | *Eulemur rufifrons* (west) | 8 | Leaves, fruits, flowers | Dry forest | Isalo |
|  | *Lemur catta* | 4 | Leaves | Dry forest | Isalo |
|  | *Varecia variegata* | 2 | Fruits | Rainforest | Ranomafana |
| Indriidae | *Avahi peyriersai* | 7 | Leaves | Rainforest | Ranomafana |
|  | *Propithecus edwardsi* | 4 | Leaves, fruits | Rainforest | Ranomafana |
|  | *Propithecus verreauxi* | 4 | Leaves, flowers | Dry forest | Isalo |

**Figure 1:**
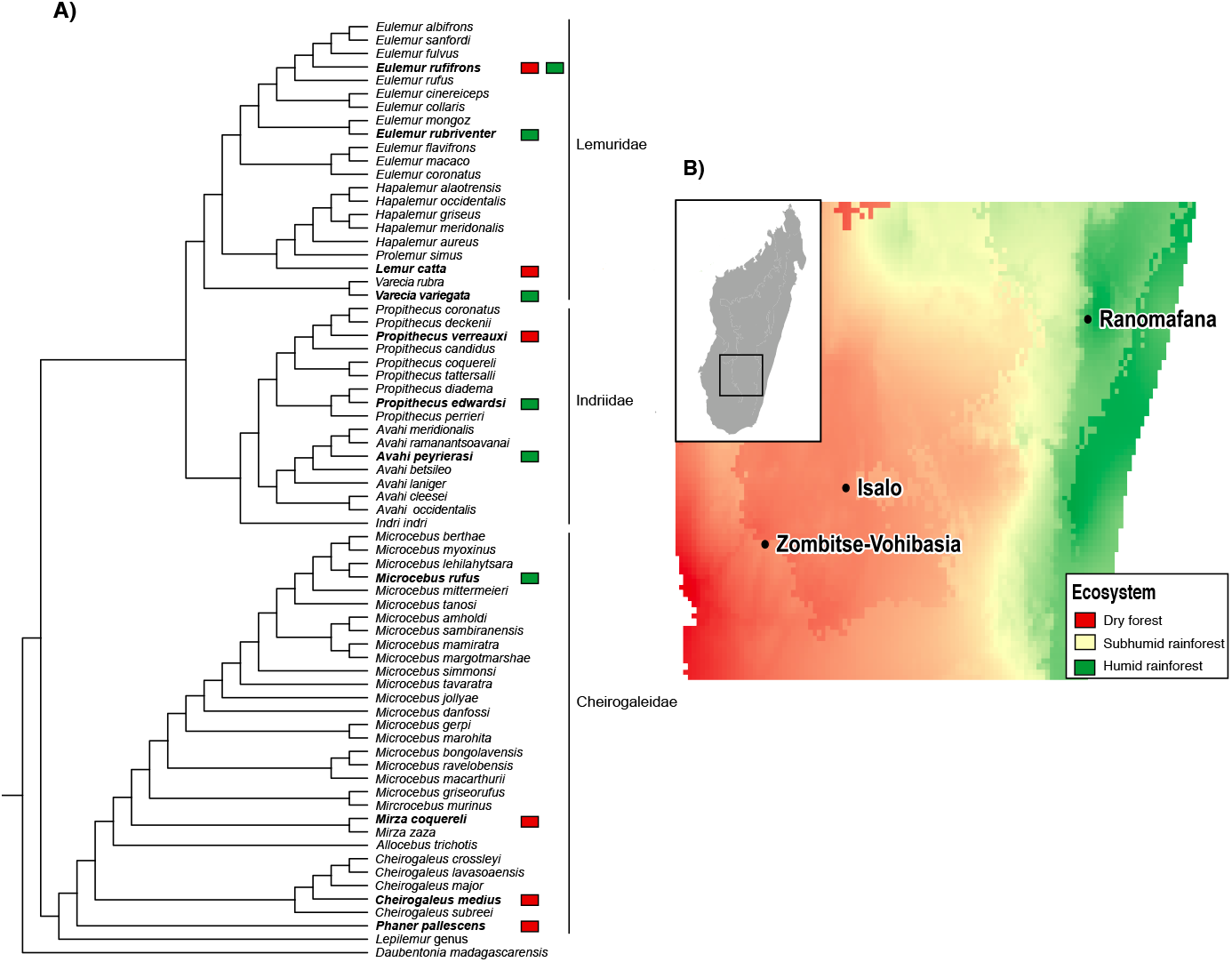
Sampling sites and species. **(a)** Lemur phylogeny adapted from Herrera & Dávalos (2016). Bolded tips denote species sampled in this study. Colors within squares represent the habitat of the collection site (see Fig. 1B). **(b)** Map of sampling sites, colored by values in the WorldClim “Precipitation of Warmest Quarter” raster layer (Hijmans et al., 2005). Figure adapted from Donohue et al. (2022).

**Figure 2:**
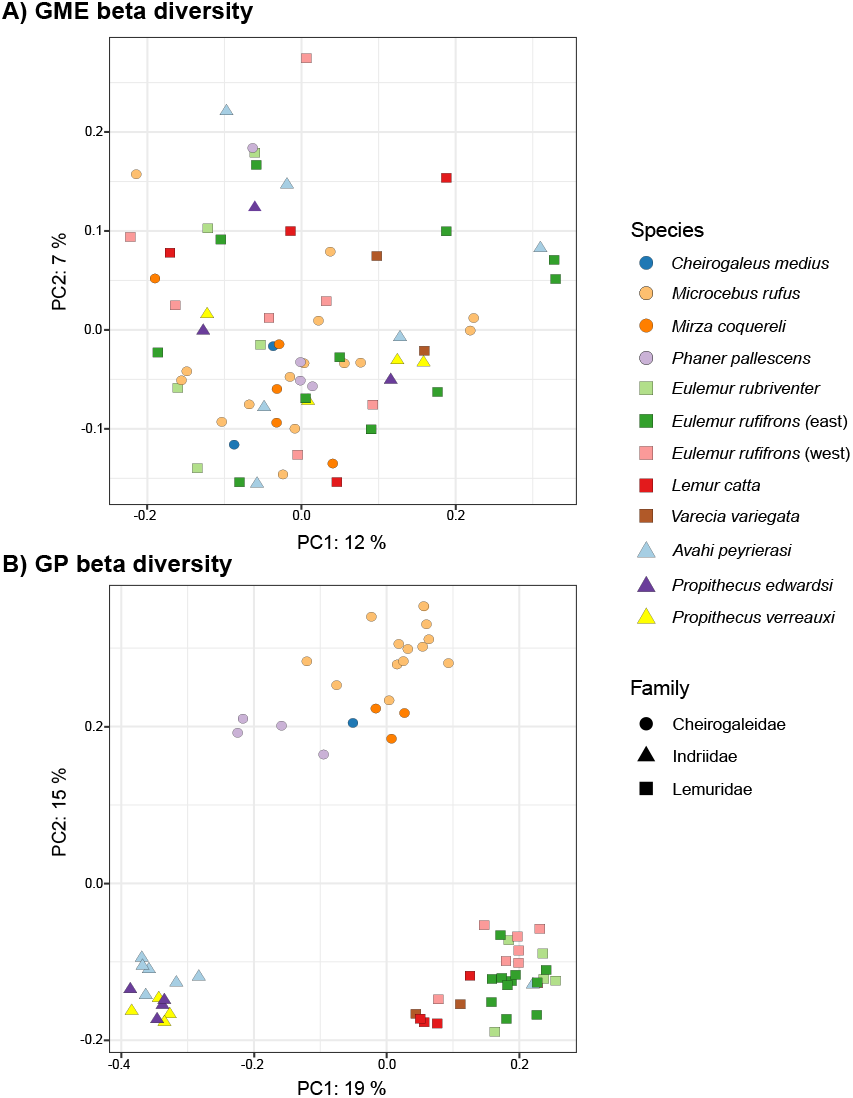
Principal coordinate analysis plots of gut microbiome unweighted UniFrac beta diversity. Colors correspond with species and shapes correspond with host family. **(a)** Ordination plot of gut microeukaryote diversity; **(b)** Ordination plot gut prokaryotic diversity.

### Data generation

For the GP portion of this study, we used 16S rRNA sequence data (v4 region) downloaded from NCBI BioProject PRJNA723621 (Donohue et al., 2022). To generate GME sequence data, we PCR amplified the V9 region of the 18S rRNA gene (Bradley et al., 2016) using preserved DNA from samples used in Donohue et al. (2022). PCRs were prepared in 20 μl volumes as follows: 10 μl of Extract-N-Amp PCR reaction mix, 4 μl of DNA template, 4 μl of mammal blocker (Vestheim & Jarman, 2008), 1 μl of H_2_O, 0.5 μl of forward primer, and 0.5 μl of reverse primer. Amplification was performed with a 3 min. denaturation period at 94ºC followed by 35 cycles of 45 s at 94ºC, 15 s at 65ºC, 30 s at 57ºC, and 90 s at 72ºC, with a final extension of 10 min. at 72ºC. To control for batch effects, DNA extractions and PCRs were randomized across species and sites. We checked the size and quality of PCR products using agarose gel electrophoresis. Libraries were normalized using the SequalPrep™ Normalization Plate Kit and quantified with a Qubit® fluorometer. After confirming each sample had a DNA concentration of approximately 1 ng/μL, samples were pooled and submitted to the UK HealthCare Genomics Core for sequencing on an Illumina MiSeq flowcell using 250 bp paired-end chemistry.

### Bioinformatics

We analyzed and re-analyzed 18S rRNA and 16S rRNA sequences, respectively, using the vsearch pipeline (Rognes et al., 2016) in QIIME2 (Bolyen et al., 2018). After clustering OTUs at 99% (for different clustering widths, see supplemental materials) and removing chimeras, we assigned taxonomy using the SILVA reference database (version 132, Quast et al., 2013). For GMEs, we also filtered obvious contaminants (i.e., bacteria, archaea, and sequences that could only be explained by diet and/or environmental exposure: Craniata, Arthropoda, Tardigrada, Myriapoda, Hexapoda, Mollusca, Archaeplastida, and Cryptophyceae; for other filtering regimes, see supplemental materials). We chose this clustering width and filtering regime because it captures the highest taxonomic diversity and includes all possible gut residents while minimizing transients (i.e., non-residents).

### Characterization of microbial species composition

As a first step towards describing the taxonomic composition of the lemur gut microbiome, we calculated the relative abundance of GMEs and GPs.

We next sought to define the core lemur microbiome. Here, the core microbiome is defined as GPs and GMEs present in at least 50% of samples. This analysis, executed using the core-features command in QIIME2, identifies microbial species shared across hosts. Shared (or “core”) microbes may be conserved by natural selection or, alternatively, are present because they are ubiquitous in the environment and incidentally ingested. We analyzed the core microbiome of the total lemur dataset, and then narrowed our analyses to scan for microbes present in all samples belonging to a given taxonomic family (i.e., all Cheirogaleidae, all Indriidae, and all Lemuridae). This narrowing allowed for the detection of clade-specific patterns, as some families may share a greater number of core microbes than others.

Next, we asked whether variation in microbial relative abundance was associated with host taxonomic (family, genus, species) and/or ecological (diet, habitat) variables. To do this, we used analysis of composition of microbiomes (ANCOM), which normalizes for sequencing depth and does not assume independence. ANCOM has two outputs: F-statistics, which measure effect size, and W-statistics, which quantify shifts in the number of reads attributed to a given OTU relative to others in the dataset. Generally, the higher the F- and W-statistic, the more significant an OTU’s differential abundance (Mandal et al., 2015). As with the core microbiome analyses, we performed ANCOMs on the total dataset and family-specific datasets.

### Characterization of microbial diversity

We calculated beta diversity using the unweighted UniFrac metric (for additional metrics, see supplemental materials). Clustering patterns were visually assessed using Principal Coordinates Analysis (PCoA) plots generated with the R packages qiime2R (version 0.99.34; Bisanz, 2018) and phyloseq (version 1.30; McMurdie & Holmes, 2013).

To compare the effects of host ecological and taxonomic factors on GP and GME beta diversity, we conducted Adonis tests in QIIME2 using beta diversity as the response variable and the formula “Diet_PC1*Habitat_PC1 + Family/Genus/Species” as the independent variable. Diet and habitat values for each species were gleaned from Donohue et al. (2022), where we conducted a literature review of previous feeding behavior studies (for Diet_PC1) and built ecological niche models based on species presence/absence data from GBIF.org (for Habitat_PC1). As these procedures output multivariate data (i.e., diet: percent of time spent feeding on fruit, leaves, insects, gum, and flowers; habitat: the 19 different WorldClim bioclimate variables (Hijman et al., 2005)) and Adonis tests require univariate data, we then performed Principal Component Analysis and extracted eigenvalues associated with PC1. Adonis tests were performed with 999 permutations using additive sums of squares. Each variable was checked for homogeneity of variance.

Next, we calculated two measures of alpha diversity for each dataset, Faith’s PD and observed OTUs, to explore the phylogenetic and taxonomic richness of each sample, respectively. Significance of associations between alpha diversity, host taxonomy, habitat, and diet were assessed using Kruskal-Wallis tests (Kruskal & Wallis, 1952) with Bonferroni corrections, and visualized with boxplots created in ggplot2.

### Phylogenetic signal of GME beta diversity

Phylogenetic signal analyses assess whether trait values in multi-species comparative datasets are conserved or divergent with respect to phylogenetic relationships. Here, we used Blomberg’s *K*, a variance ratio that measures whether closely-related species resemble each other more (*K* > 1) or less (*K* < 1) than expected under Brownian motion models of evolution (random drift; *K* = 1) (Blomberg et al., 2007). Because Blomberg’s *K* requires a single trait value for each tip, we built three subsampled datasets with randomly selected samples representing each species. Because Blomberg’s *K* also requires univariate data, we performed a PCoA of beta diversity and tested phylogenetic signal of the first principal coordinate (PCo1). Finally, we explored whether the presence of low-abundance OTUs weakens phylogenetic signal by filtering the three subsampled datasets and comparing results; the “unfiltered” dataset includes all OTUs, the “> 100” dataset excludes OTUs with fewer than 100 reads, and the “> 1000” dataset excludes OTUs with fewer than 1,000 reads. Blomberg’s *K* was calculated using the “phylosig” function in the R package Geiger (version 2.0.7; Harmon et al., 2007). The lemur phylogeny was built using a pruned Bayesian fossilized birth death tree published in Herrera & Dávalos (2016), downloaded from Dryad (https://datadryad.org/stash/dataset/doi:10.5061/dryad.51f00). Here, we only analyze GME phylogenetic signal, as GP results were reported in Donohue et al. (2022).

### Cross-kingdom interactions

To better understand cross-kingdom interactions within the lemur gut, we first asked whether a relationship existed between the species richness of GP and GME communities. To do this, we conducted Pearson correlation tests on GP and GME alpha diversity (observed OTUs) for each sample.

Next, we used undirected co-occurrence network analysis to identify significant interactions between GP and GME OTUs. We removed all OTUs with fewer than 1,000 reads to reduce noise and complexity. To find OTUs with significant co-occurrence patterns (i.e., relative abundance of OTU1 is correlated with the relative abundance of OTU2), we performed Spearman’s rank correlation analyses with the R package Hmisc (version 4.4; Harrell, 2020).

We then visualized significant interactions (Spearman p-value < 0.05) using Cytoscape (Shannon et al., 2003), setting the p-values and correlation coefficients (r-values) as edge attributes. We generated two networks, one that included all significant microbe-microbe interactions, and another that only included significant cross-kingdom interactions. Network summary statistics were computed using the Cytoscape “Analyze Network” function. These summary statistics (namely, centrality measures) allowed us to test the influence of GMEs on GPs. Using Wilcoxon tests, we asked whether (1) GMEs or GPs had greater connectivity, and (2) GPs connected to GMEs had greater connectivity than GPs that were not.

## RESULTS

### Microbial species composition

A total of 622,811 sequence reads (mean = 33,839 per sample) were used for the 16S rRNA GP dataset. For GMEs, we recovered 1,261,835 reads (mean = 18,026 per sample). We found that Stramenopiles, including Blastocystis, were the most abundant GMEs (22.51% across dataset), followed by Ascomycota fungi (20.82%), Alveolata protists (20.75%), Chromadorea nematodes (14.76%), Basidiomycota fungi (8.38%), Parabasalia protists (7.56%), and Cestoda platyhelminths (2.07%) (Fig. S1a). In the GP dataset, Bacteroidetes was the most common phylum (30.91%), followed by Firmicutes (27.76*%*), Proteobacteria (15.91%), Actinobacteria (6.40%), Spirochaetes (5.99%), Cyanobacteria (5.35%), and Kiritimatiellaeota (2.29%) (Fig. S1b).

Core gut microbiome analyses showed 25 GPs and 212 GMEs were shared across at least 50% of samples. Three GME OTUs were present in all samples, but none could be classified below the domain-level. Ten GMEs were present in all Cheirogaleidae samples, four of which could be classified: Blastocystis, Chromadorea nematodes, *Papiliotrema terrestris* strain LS28 (a Basidiomycota fungus), and an unknown *Bradymyces* species (an Ascomycota fungi). Of the nine core Indriidae GMEs, just three were classified: Chromadorea nematodes, the same unknown *Bradymyces* species as in Cheirogaleidae, and *Paraschneideria metamorphosa* (a species of Alveolate protist). No GME OTUs were shared among all Lemuridae. Core gut microbiome analyses also showed no GP OTUs were shared within Cheirogaleidae, Indriidae, or across all samples. Two Firmicutes, however, were shared among Lemuridae: Erysiplelotrichaceae and Lachnospiraceae.

ANCOM analysis did not identify differentially abundant GMEs associated with host diet, ecosystem, family, or genus. It did show, however, that an Echinamoebida Amoebozoa was more abundant in *Cheirogaleus medius* than any other species (W = 55; F = 5). In contrast, many GPs were associated with host taxonomy, diet, and habitat (supplemental results).

### Microbial diversity

PCoA plots indicated that while GP beta diversity partitioned according to host family, GMEs did not cluster in association with any study variable (Fig. 3; Figs. S2-S6). Adonis tests showed that diet, habitat, and host taxonomy had no effect on GME beta diversity, whereas associations with GPs were consistently significant. GME alpha diversity did not differ across study variables; GP alpha diversity was significantly associated with diet and host taxonomy, but not habitat (Fig. 4a; Table S2).

**Figure 3:**
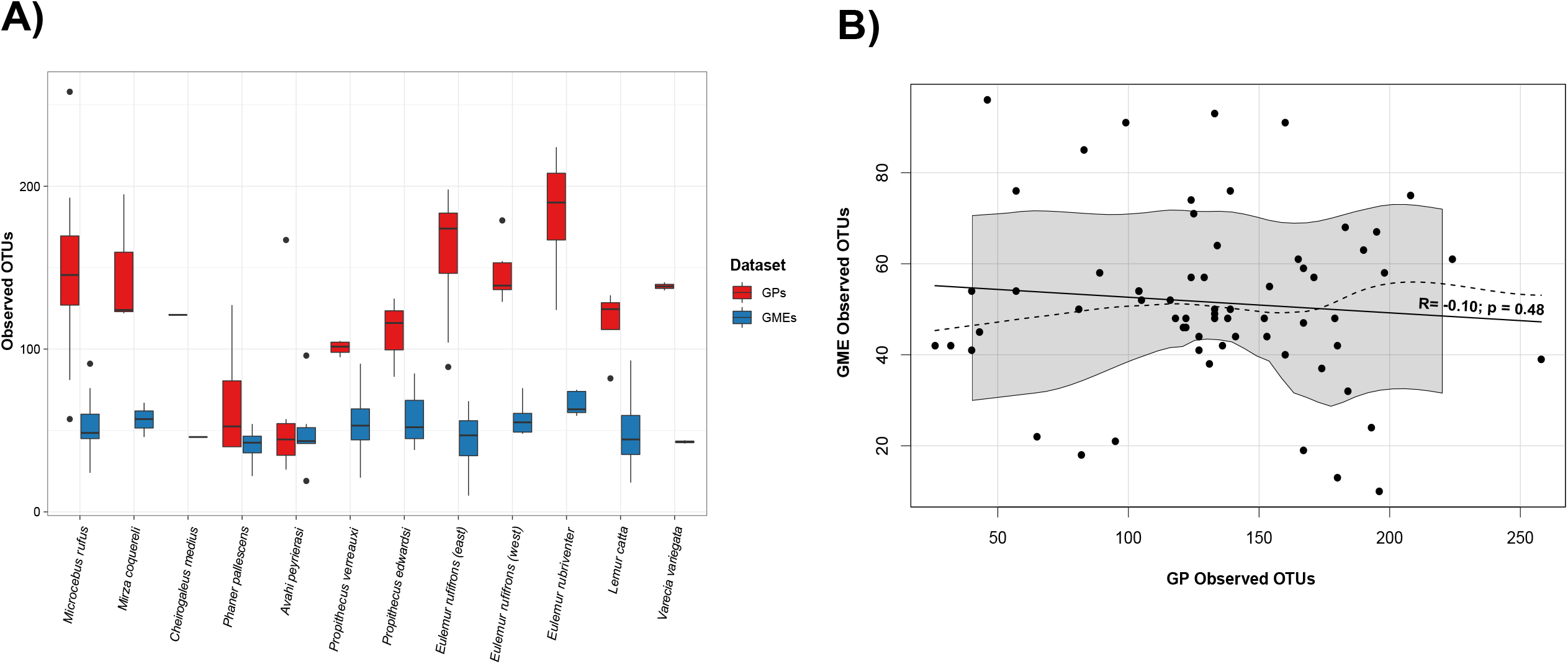
Measures of gut microbiome alpha diversity for gut microeukaryotes (GMEs) and gut prokaryotes (GPs). **(a)** Box plot shows the number of observed OTUs in GP (red boxes) and GME (blue boxes) communities. **(b)** Scatterplot showing relationship between the number of observed GP and GME OTUs for each host sample in the data set. The solid black line represents a fitted regression line, with its correlation coefficient and significance. The dashed line and shading are nonparametric estimates of the mean (loess line) and variance, respectively.

**Figure 4:**
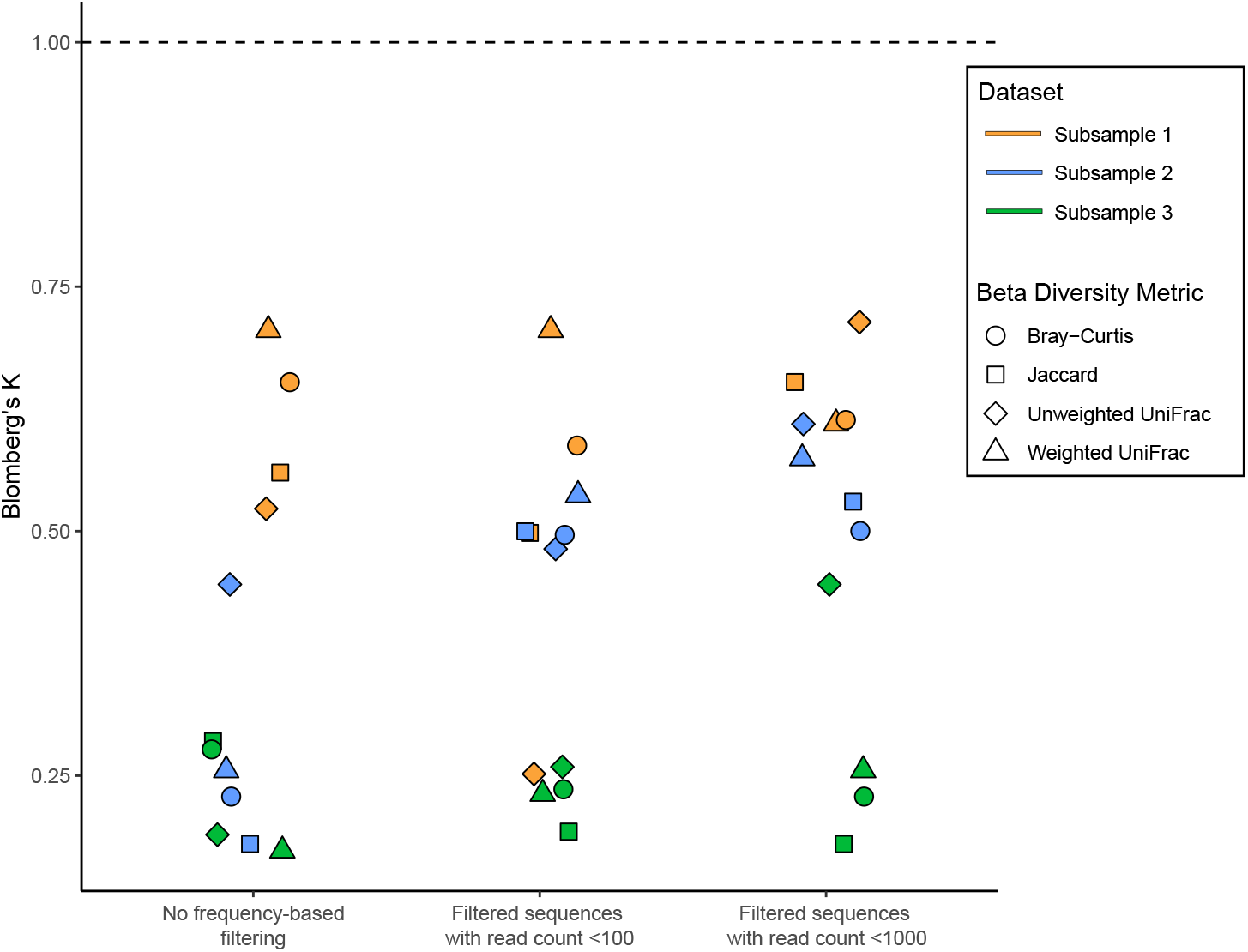
Tests of phylogenetic signal of gut microeukaryote (GME) communities as measured with Blomberg’s *K*. Tests were performed using multiple Beta diversity measures (shapes) and were applied across different OTU frequency-based filtering regimes (x-axis) and host-tip subsampling replicates (colors). All measures of *K* were non-significant (p > 0.05), indicating no phylogenetic signal of GME communities across lemur host phylogeny.

### Phylogenetic signal of GME beta diversity

Lemur GME communities were more divergent than expected under Brownian motion models of trait evolution (Fig. 5). All tests of phylogenetic signal using Blomberg’s *K* were insignificant (p > 0.05), consistent with random assemblages of GMEs in lemurs. All *K* values were lower than one (average = 0.43). ANOVA tests showed *K* values did not differ across beta diversity metrics (p = 0.95) or filtering regimes (p = 0.27). There were significant differences across subsampled datasets (p < 0.01), with average *K* values of 0.59, 0.25, and 0.44 in subsampled datasets 1-3 respectively.

**Figure 5:**
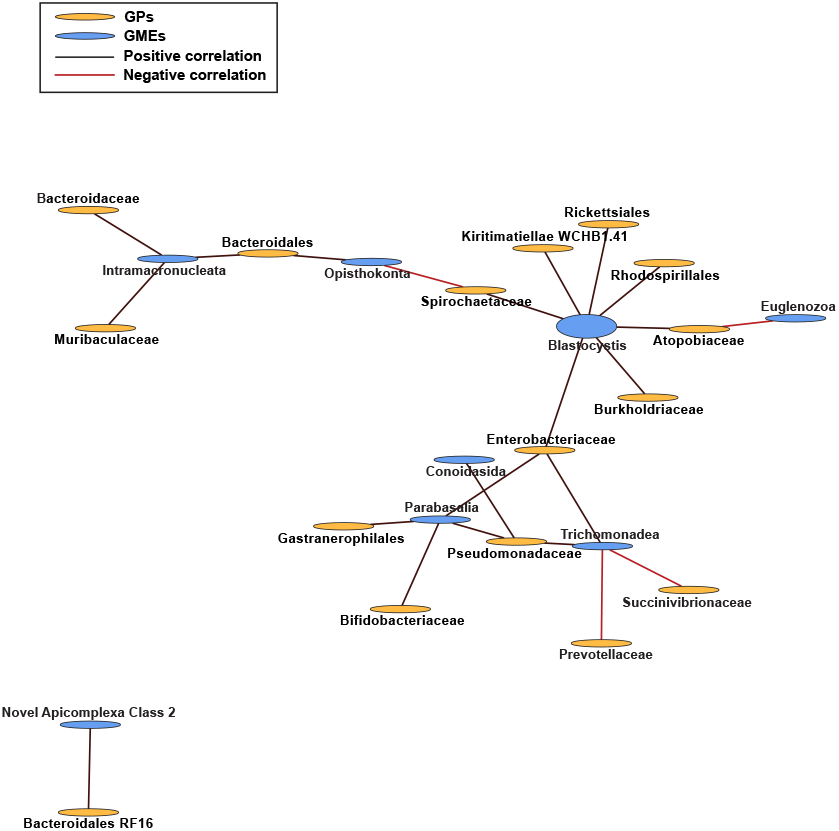
Microbial co-occurrence network analysis identifying significant interactions between gut prokaryote OTUs (orange ovals) and gut microeukaryote OTUs (blue ovals). Black lines indicate positive correlations (e.g., increased abundance of Pseudomonadaceae was associated with increased abundances of Trichomonadea and Parabasalia), while red lines indicate negative correlations (e.g., increased abundance of Trichomonadea was associated with a decrease in Prevotellaceae and Succinivibrionaceaee). Oval size increases with centrality score.

### Cross-kingdom interactions

We did not detect a significant relationship between GP and GME species richness (R = -0.10; p = 0.48) (Fig. 4b).

In the full-dataset cross-kingdom network, we found that of 1,800 pairwise comparisons, read counts between OTUs were significantly correlated in 671 (37%). GME nodes had significantly lower closeness centrality scores than GPs (p < 0.001), but their betweenness centralities were comparable (p > 0.05). Focusing on the strictly cross-kingdom network (i.e., only includes connections between GPs and GMEs, not GP-GP or GME-GME), we detected 24 significant correlations (Fig. 6). GPs that were significantly correlated with GMEs did not have higher centrality scores than GPs that were not (p > 0.05). Blastocystis was the most connected OTU (closeness centrality score = 0.42) and had the greatest number of significant correlations (7 of 24) (Fig. 6). Blastocystis read count was positively correlated with Enterobacteriaceae (p = 0.01; R = 0.32), Atopobiaceae (p = 0.01; R = 0.32), Rhodospirillales (p = 0.03; R = 0.28), Kirimatiellae WCHB1.41 (p = 0.03; R = 0.28), Spirochaetaceae (p = 0.03; R = 0.28), Burkholderiaceae (p = 0.04; R = 0.26), and Rickettsiales (p = 0.04; R = 0.26).

## DISCUSSION

This is one of the first studies to explicitly compare diversity patterns between GP and GME communities. This type of analysis is useful because it offers a more comprehensive picture of gut microbiome structure, diversity, and evolution, with new insight into how different microbial communities shape and are shaped by host ecology and evolution. Surprisingly, we found that while lemur GPs vary with host taxonomy, diet, and habitat, GMEs have no detectable association with any of these factors. This striking difference between different components of sympatric microbial communities raises many new questions as to how the gut microbiome evolves and influences host biology.

### Contrasting patterns between prokaryotic and microeukaryotic communities

The observed differences in diversity patterning between GPs and GMEs suggest these communities serve fundamentally different roles and are therefore shaped by different forces. From these results, we posit GPs represent natural extensions of the host, inextricably linked to survival, fitness, and homeostasis, while GMEs, as a community, primarily act as foreign bodies.

GPs have formed stable relationships with hosts and reflect millions of years of coevolution and ecological filtering based on phylogenetically structured traits, including feeding strategy and gut morphology (Amato et al., 2019). Intraspecific variation in GPs, especially within populations, is generally relatively low, though factors such as sociality, seasonality, and age can lead to significant differences (McKenney et al., 2015; Bennett et al., 2016; Perofsky et al., 2017; Raulo et al., 2018; Perofsky et al., 2021; Murillo et al., 2022). Low GP variability within species largely stems from natural selection; purifying selection purges microbes with negative effects, while positive selection promotes the association between host and microbes that yield a benefit, together driving the adaptive evolution of endogenous GP communities (Garud et al., 2018). Such strong selective pressure through evolutionary time likely arises from the crucial roles GPs play in host immunity and nutrient acquisition. Different host species have evolved different requirements to maintain homeostasis, leading to unique GP communities. As demonstrated by reciprocal transplant studies, hosts suffer severe reductions in fitness without their species-specific GPs – even if they are implanted with the GPs of closely-related species (e.g., Brooks et al., 2016; Moeller et al., 2019; van Opstal & Bordenstein, 2019; Parker et al., 2021).

Our evidence from a comparative approach in lemurs showed no clear links with GME diversity and ecological or evolutionary variables, leading to the conclusion that GME communities perform few functions promoting host homeostasis and are therefore not maintained by selection. If, like GPs, GMEs as a community were crucial to host functioning, we would expect to find signatures of phylogenetic signal or adaptation. However, GME communities were largely random (Fig. 3). Furthermore, phylogenetic signal analyses confirmed no correlation between GME diversity and host evolutionary relationships (Fig. 5). In contrast, alpha diversity was surprisingly constrained across samples (Fig. 4), which suggests high GME species richness may negatively impact host fitness. It is possible the host immune system treats most GMEs as pathogens, even though many taxa act as true symbionts and interact with biologically important GPs (e.g., Blastocystis; Fig. 6). As an explanation, we suggest that lemur GMEs comprise a greater proportion of both commensal and parasitic taxa compared with GPs; because the immune system removes taxa producing a negative effect, far fewer taxa develop positive long-term associations with a host. Unfortunately, we cannot test this hypothesis because it is not yet possible to distinguish parasitic from non-parasitic microbes molecularly; instead of innovating new traits specific to parasitism, it seems parasites have simply modified existing traits conserved across microbes, rendering them genetically invisible (del Campo et al., 2020).

### Are lemur GME communities exceptionally random?

It is unclear whether our contrasting GP and GME results are unique to lemurs or can be extrapolated across the Tree of Life. One possibility is that the ecology and behavior of lemurs precludes an abundance of resident GMEs. This was first hypothesized in Mann et al. (2020), which, to our knowledge, is the only other study (not just in Primates) to examine GMEs in a phylogenetic context, making comparisons across lemurs, catarrhines, platyrrhines, and apes.

We believe there are two comparisons to be made between our study and that of Mann et al. (2020). First, it’s important to consider the issue of scale, both phylogenetically and geographically. Broad phylogenetic sampling of host species has been previously shown to amplify gut microbiome phylogenetic signal, whereas narrowing the dataset to more closely-related species tends to reduce phylogenetic signal and amplify ecological effects (Greene et al., 2019). This difference in scale may help explain why Mann et al. (2020), which sampled broadly across the Primate order, detected phylogenetic signal of GME communities while we did not. A meta-analysis would help test the effects of scale; unfortunately, however, we amplified a gene region that was not sequenced in Mann et al. (2020). Another possibility is that GME communities are truly random with respect to phylogeny, and that the phylogenetic effects detected in Mann et al. (2020) reflect the wide geographical spread of the species they examined. Different ecosystems and continents have different microeukaryotic assemblages, presenting the opportunity for incidental ingestion of taxa endemic to the host’s extrinsic environment. To help disentangle the effects of phylogenetic and geographic scale, future efforts should prioritize comparisons among sympatric and phylogenetically diverse species.

Second, it is important to consider the possibility that lemurs harbor exceptionally random GME assemblages. This idea is supported by Mann et al. (2020), which found *Propithecus verreauxi* to have the fewest GME taxa. It was reasoned that because *Propithecus verreauxi* is almost entirely arboreal and obtains water from food, this species has limited opportunity to contract eukaryotic organisms that are transmitted through feces, soil, and freshwater. Extrapolating across our dataset, arboreality is true of nearly every species and may explain the lack of an effect of habitat on GME patterning despite sampling lemurs in vastly different ecosystems (Fig. S3-S6).

However, if contact with soil is indeed a major force shaping GME communities, we would expect significant differences between *Lemur catta* and all other species. *Lemur catta* is the most terrestrial lemur species and is known to practice geophagy and drink cave water in Isalo National Park (Sauther et al., 2013), where our samples originated. If terrestriality were a major factor, we would expect to find less intraspecific variation in GME composition and diversity within *L. catta* compared with all other species. We might also expect to find significant differences in the presence/absence and/or abundance of microeukaryotic indicator species – for example, higher abundances of taxa found in soil. However, we did not find any such differences between GME patterns of *Lemur catta* and other, more arboreal lemurs.

### Resident and transient members of the lemur eukaryome

Of the GMEs identified in our data, it remains unclear which taxa constitute “true” members of the lemur gut microbiome. Here, we define “true” as resident inhabitants of the lemur gut. However, differentiating resident and transient GMEs is difficult without longitudinal sampling, so we can only make assumptions based on previous research. If not a “true” member of the host’s gut microbiome, species may represent a dietary component, pathogen, or environmental contaminant. In total, we identified six major groups of microeukaryotes: alveolates, stramenopiles, parabasalians, chromadorea nematodes, cestodes, and fungi. Due to low taxonomic resolution, it is not yet possible to characterize the relationship of GMEs identified via metabarcoding, as microbes within these groups exhibit a range of symbioses. Nonetheless, we can speculate.

Transient members of the eukaryome are typically acquired from the host’s diet and environment. Interestingly, we found a high proportion of microbes classified as arthropod symbionts – namely, gregarines and parabasalians (Adl et al., 2012) – which likely entered the lemur gut through insectivory. This result is not surprising considering Rowe et al. (2021)’s discovery of insect DNA in the feces of lemur species never before observed consuming animal prey, including indriids, a primarily folivorus taxa. Fungal sequences presented a greater challenge, as they can stem from dietary components and obligate symbioses. While generally uncommon in primates, lemurs do practice both opportunistic and accidental mycophagy (Hanson et al., 2003; Borruso et al., 2021). Although we also detected possible environmental microbes, including CONthreeP alveolates and free-living worms that could inhabit soil and water (Fierer, 2017; Fernandes et al., 2021), these likely account for a smaller proportion of transients than diet. We reached this conclusion after finding no differences between GMEs collected from Madagascar’s eastern rainforests and western dry forests – two vastly different habitats which certainly harbor unique microbial assemblages.

Pathogens occupy a classification between transient and resident, as they can specialize and propagate on a host for extended periods of time. We recovered several possible pathogens, including litostomes and worms. Though some litostomes are known commensals, many serve as enteric parasites that have been reported in primates (Adl et al., 2012; Parfrey et al., 2014). We also identified nematodes (Chromadorea) and cestodes (i.e., tapeworms) in our data, both of which represent a variety of parasitic worms (Aivelo et al., 2015).

Finally, we identified many candidate resident GMEs. We detected highly abundant mammalian commensals in our dataset, including the protists litostomatea, parabasalians, and blastocystis. Blastocystis are particularly interesting, as they have been associated with both healthy and diseased gastrointestinal tracts in humans (Deng et al., 2021). In our network analyses, blastocystis were the most centrally connected GME taxa, which suggests an important role in shaping cross-kingdom interactions in the lemur gut (Fig. 6). Among fungi, Basidiomycota and Ascomycota were present in nearly all samples and have already been described as dominate phyla in the primate gut (Sharma et al., 2022). We also found that commensalism may not necessarily be a pre-requisite of gut “residency”, as parasitic Chromadorea nematode reads were nearly ubiquitous.

### Conclusions and future directions

This study represents an important first step towards understanding GME community structure and diversity in primates. We conclude that, when analyzed on a phylogenetic scale, GMEs are not shaped by host evolution, diet, or habitat. However, it is important to note that other lemur studies detected effects of seasonality on GME communities (Aivelo et al., 2015; Murillo et al., 2022). These studies differed from ours, as they re-sampled the same populations at multiple time-points and focused on a single host species. Relatedly, many microscopy studies have shown lemur endoparasite composition to shift with ecological factors over time (e.g., Raharivololona & Ganzhorn, 2010; Benavides et al., 2012; Springer & Kappeler, 2016). Therefore, it is possible that seasonality is a predominant force shaping GME communities – a factor we could not test given our study design. Multi-species longitudinal studies may reveal lemur GMEs are less random than the evidence we presented herein.

In addition to emphasizing the insight we could gain from more GME metabarcoding surveys at different taxonomic and temporal scales, we also highlight the need for more complete genetic databases. In publicly available databases, 80% of sequences belong to bacteria and 20% to eukaryotes, including plants and animals (del Campo et al., 2020). Without the ability to identify GMEs to the species-level, it is impossible to classify their relationship with the host. This context is critical, as we hypothesize commensals and parasites within the eukaryome are shaped by different factors and should be analyzed as distinct communities. Database expansion would also facilitate more reliable and rapid assessments of parasite infection in endangered species.

Though this study created more questions than it answered, it, at the very least, offers a compelling argument to shift the paradigm and vocabulary of gut microbiome research. The gut microbiome is comprised of many “omes” (i.e., communities), all of which are structured according to different forces with “ome”-specific consequences – at least in lemurs. Future work should always clarify the target community with appropriate data, whether it be the prokaryome, eukaryome, mycobiome, virome, or the entire microbiome.

## Supporting information

Supplemental matrials

## DATA ACCESIBILITY

Data accessibility: 18S rRNA raw reads have been uploaded to the NCBI database (BioProject PRJNA913825) and will be released upon manuscript acceptance. 16S rRNA reads were published under NCBI BioProject PRJNA72362.

## ACKNOWLEDGEMENTS

Methods for fecal collections of wild primates were approved by the IACUC committees of the University of Kentucky (IACUC #2018-2919) and Stony Brook University (IACUC #1177457-3 and #11323621-2). Field protocols were approved by Madagascar National Parks and by the Ministere de l’Environment et des Eaux et Forets.

This work was supported by the following funders: the Animal Behaviour Society Student Research Grant; the American Society of Primatologists General Small Grant; the Global Wildlife Conservation’s Lemur Conservation Action Fund and IUCN’s Save Our Species (SOS); Grant-in-Aid of Research from Sigma Xi, The Scientific Research Society; HHMI Sustaining Excellence-2014 grant (#52008116), Primate Conservation, Inc.; Rowe-Wright Primate Fund; Society of Systematic Biologists Graduate Research Award; a Stony Brook University Graduate Student Employment Union Professional Development Award; a Stony Brook University Interdepartmental Doctoral Program in Anthropological Sciences Research Award; and the University of Kentucky Ribble Endowment Fund.

We thank the Madagascar Ministry of Environment, Forests, and Ecology; and Madagascar National Parks in Andringitra National Park, Isalo National Park, Ranomafana National Park, and Zombitse-Vohibasia National Park for allowing us to conduct this field research. We thank Centre ValBio research station and MICET/ICTE for providing logistical support in Madagascar.

