## Supplemental matrials for "Lemur gut microeukaryotic community variation is not associated with host phylogeny, diet, or habitat"

#### Supplementary Methods

##### Characterization of microbial diversity sensitivity analyses

For microbial diversity analyses, we compared results across different OTU clustering widths (90%, 94%, 97%, and 99% similarity thresholds) to assess whether community partitioning primarily reflects nascent (i.e., 99% clustering) or ancient (i.e., 90% clustering) microbial evolution, in addition to discerning whether community partitioning is strongest at higher or lower taxonomic levels (90% and 99% clustering, respectively). All datasets were rarefied to account for differences in sequencing depth.

From there, we conducted three different GME filtering regimes: (1) no filtering (i.e., all microbes included), (2) filtering obvious contaminants, (3) filtering obvious contaminants plus fungi, as it remains difficult to differentiate true members of the primate gut mycobiome and dietary components (Lai et al., 2019). We conducted this filtering sensitivity analysis to determine whether statistical associations between GME community diversity and study variables were strengthened once contaminants were removed.

We calculated four measures of beta diversity (unweighted UniFrac, weighted UniFrac, Bray-Curtis, and Jaccard) for each dataset to examine how differences in microbial species composition arose across samples. Leveraging different metrics can help illuminate how microbial communities vary; UniFrac metrics emphasize the effects of older microbial clades while Bray-Curtis and Jaccard are more sensitive to recent microbial evolution (Sanders et al., 2014; Donohue et al., 2022). Further, comparing weighted (weighted UniFrac and Bray-Curtis) and unweighted (unweighted UniFrac and Jaccard) metrics can indicate whether shifts are primarily driven by differences in the relative abundance of shared microbes or the presence/absence of distinct microbes across samples, respectively.

#### RESULTS

For the 18S rRNA GME dataset, the number of sequence reads varied across filtering regimes. We recovered 1,719,490 reads in the unfiltered dataset (mean = 24,565 per sample), 1,261,835 reads in the dataset with obvious contaminants filtered (mean = 18,026 per sample), and 1,046,819 reads in the dataset with obvious contaminants and fungi filtered (mean = 14,955 per sample).

##### GP ANCOM

Scanning across the entire dataset, the GPs of *Microcebus* and *Mirza* – two cheirogaleid genera with diets mainly comprising fruits and insects – harbored elevated abundances of Fusobacterium (W = 919, F = 200). *Propithecus* had higher abundances of Delfuviitaleaceae UCG-011 (W = 908; W = 110). Lemurs in dry forest habitats had more Anaerorplasma (W = 916, F = 2.5).

Comparisons within families revealed that *Cheirogaleus medius* has significantly more *Candidatus methanogranum* than other cheirogaleids. The Indriidae-specific dataset showed that *Propithecus verreauxi* (our only dry forest Indriid) had significantly more Gastranaerophilales (W = 174; F = 3.0), Succinatimonas (W = 171; F = 3.5), and Tyzzerella (W = 164; F = 2.8) than other Indriids. The *Propithecus* genus also had more Muribaculaceae uncultured bacterium (W = 223; F = 6), Lachnospiraceae NK3A20 group (W = 201; F = 5), Bacteroidales (W = 197; F = 4.5), and Fibrobacter (W = 191; F = 4.3) than *Avahi.* The Lemuridae-specific dataset showed that rainforest species had significantly more Rhodospirillales uncultured bacterium (W = 692; F = 3.0) than those in the dry forest. *Varecia variegata* had more Pasteurella (W = 645, F = 65) and Spirochaetaceae (W = 736; F = 40), *Lemur catta* had more Methanobrevibacter (W = 683; F =

17), and the *Eulmeur* genus had more Ruminiclostridium (W = 726; F = 23) and Kiritimatiellaea type WCHB1-41 (W = 745; F = 20).

### Supplementary Figures and Tables

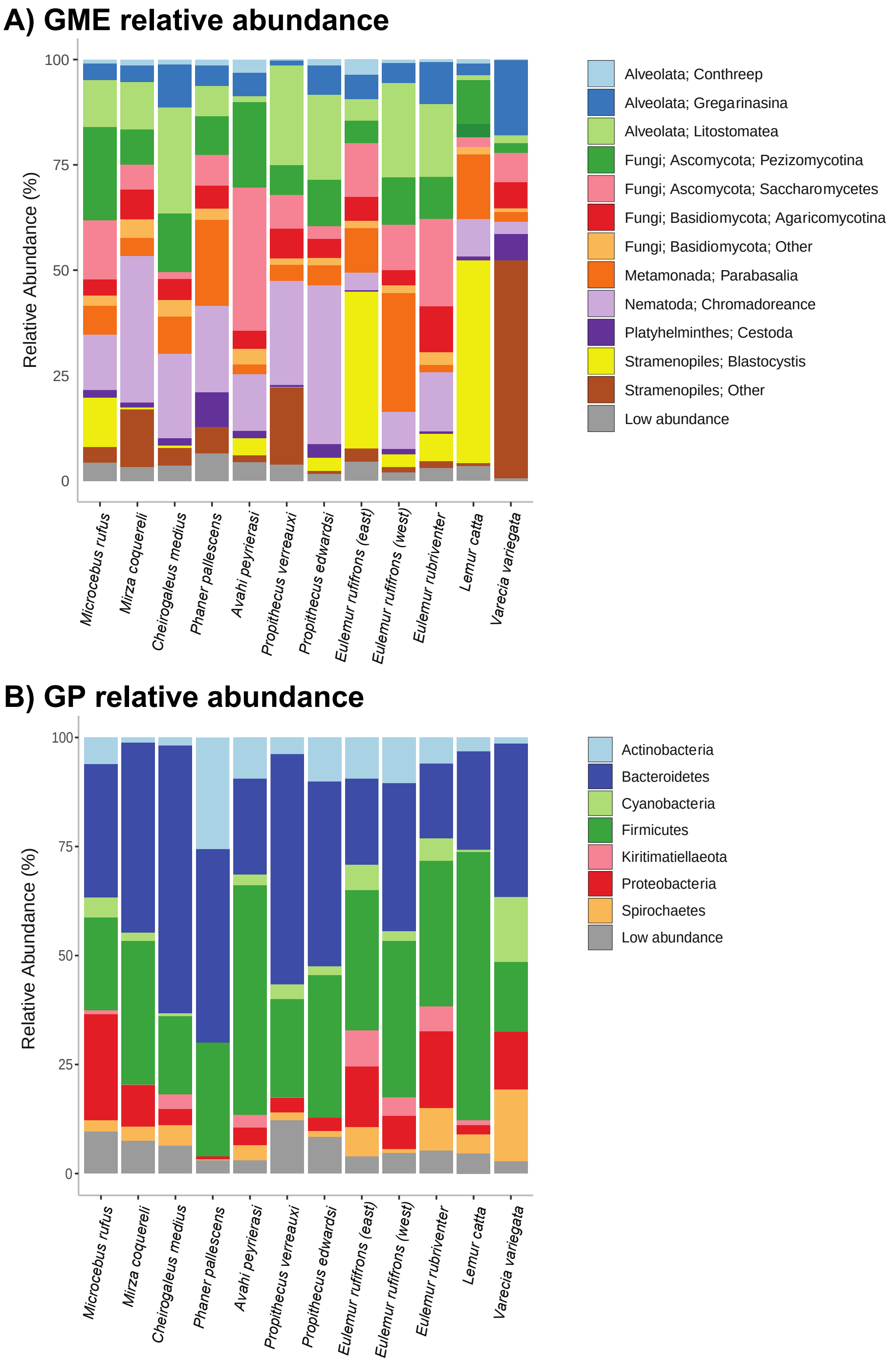


**Figure S1:** Relative abundance of major microbial taxa in lemur gut microbiomes. **(a)** Gut microeukaryote abundance at various levels of higher-level classification; **(b)** Gut prokaryote abundance at the phylum level.

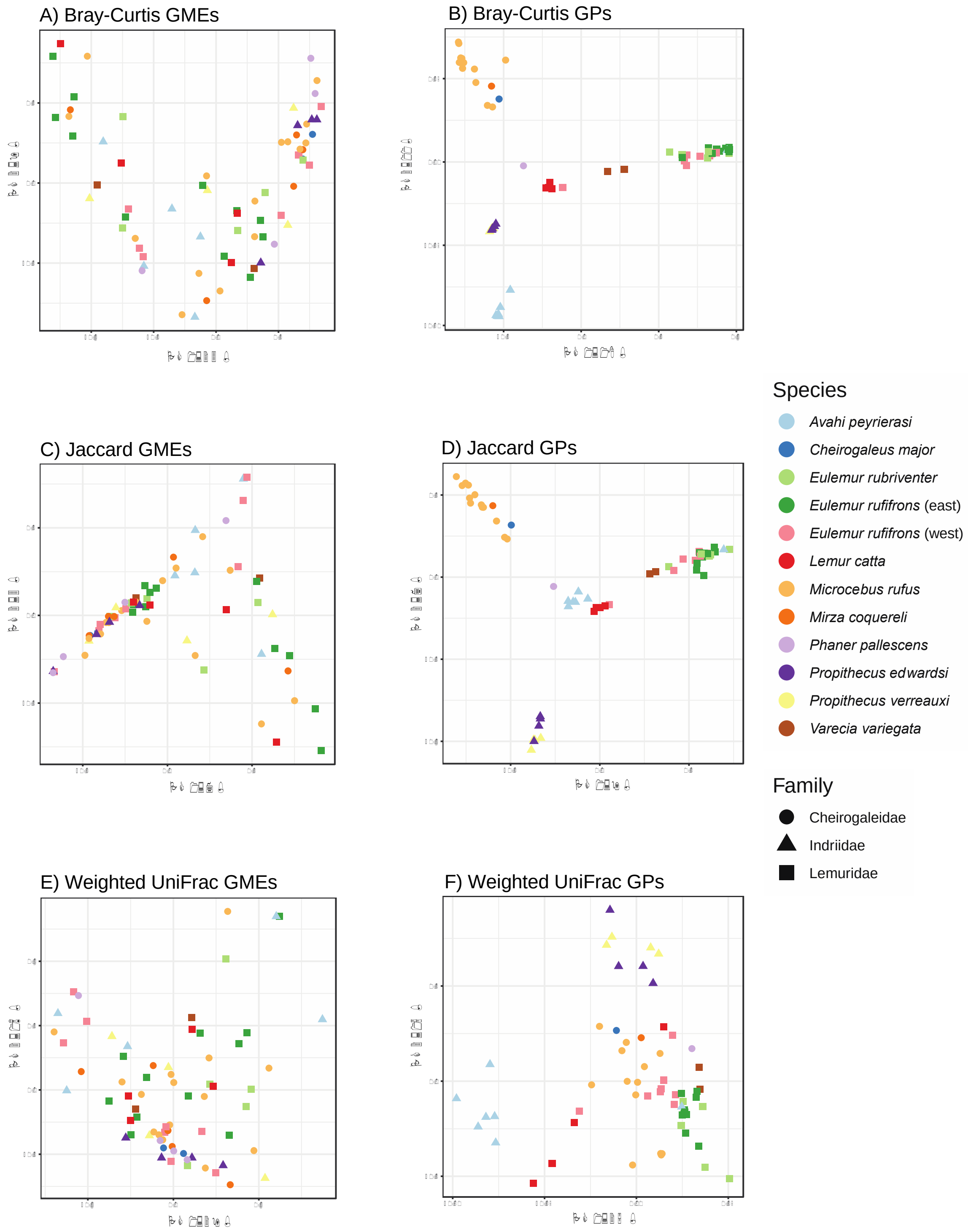


**Fig S2:** PCoA plots of gut microbiome beta diversity (unweighted UniFrac shown in the main text). Colors correspond with species and shapes correspond with family. Generally, GPs clustered according to taxonomy (right column) while GMEs did not (left column).

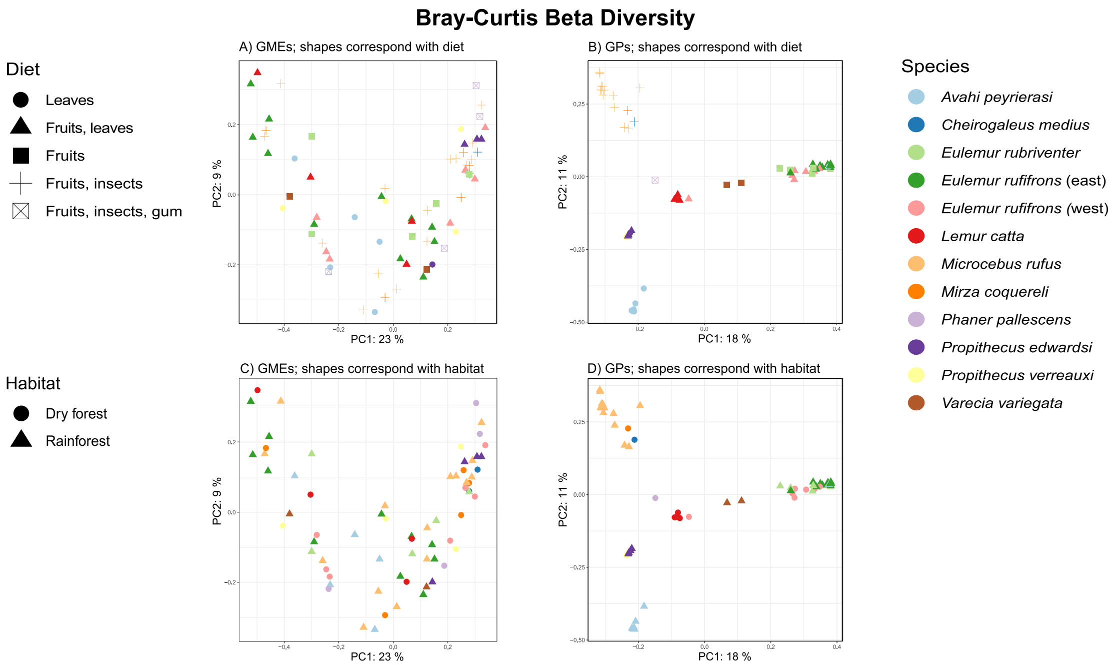


**Fig S3:** PCoA plots of Bray-Curtis beta diversity, with colors and shapes corresponding with diet and habitat, respectively. Diet, but not habitat, may contribute to GP patterning; neither factor influenced GMEs.

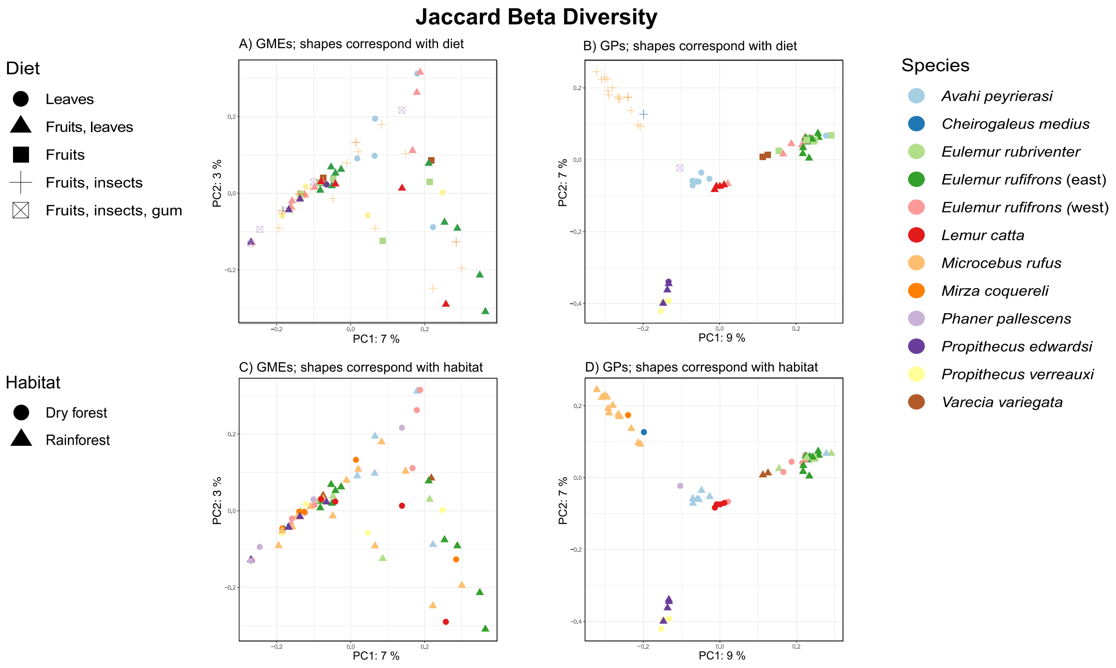


**Fig S4:** PCoA plots of Jaccard beta diversity, with colors and shapes corresponding with diet and habitat, respectively. Diet, but not habitat, may contribute to GP patterning; neither factor influenced GMEs.

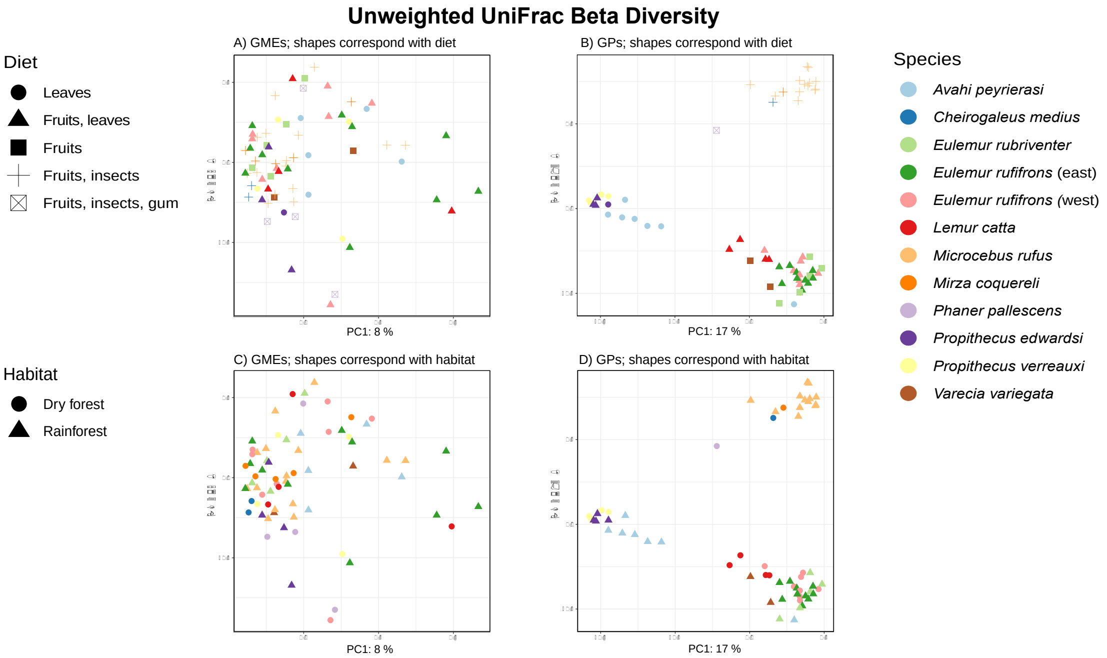


**Fig S5:** PCoA plots of Unweighted UniFrac beta diversity, with colors and shapes corresponding with diet and habitat, respectively. Diet, but not habitat, may contribute to GP patterning; neither factor influenced GMEs.

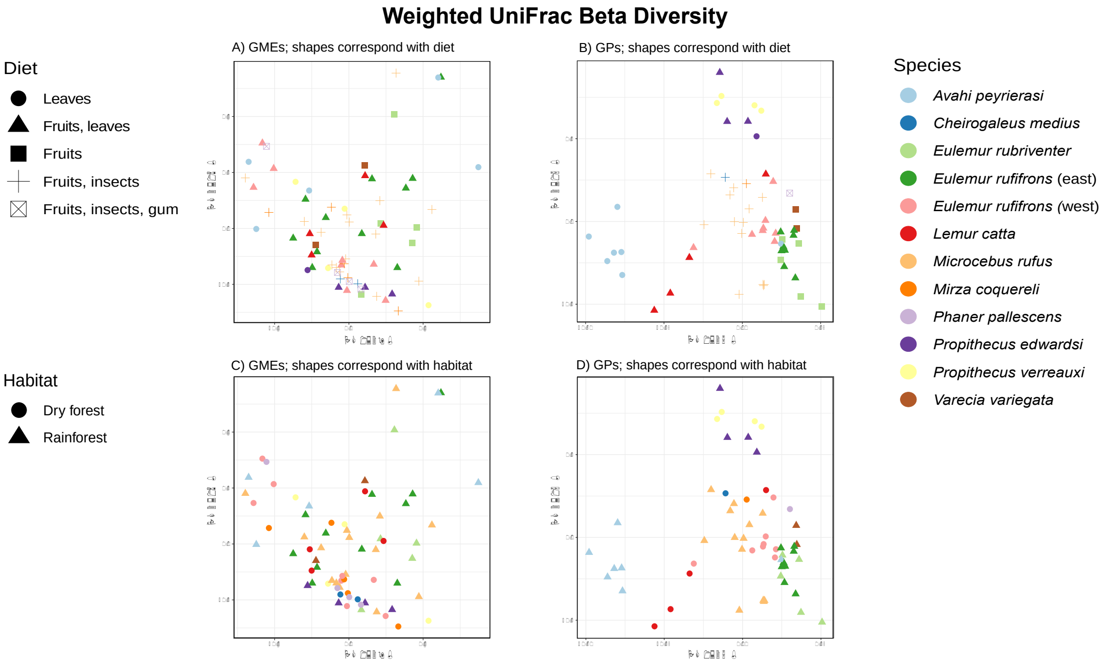


**Fig S6:** PCoA plots of Weighted UniFrac beta diversity, with colors and shapes corresponding with diet and habitat, respectively. Diet, but not habitat, may contribute to GP patterning; neither factor influenced GMEs.

**Table S1:** Adonis results for GME and GP communities. GP beta diversity was associated with host taxonomic and ecological variables, while GME beta diversity was not. GP associations tended to decrease as the clustering threshold increased.

| Community | Filter | Metric | Clustering | Variable | Adonis Test | |
|---|---|---|---|---|---|---|
| | | | | | $R^2$ | P-value |
| **GME** | Full | Bray-Curtis | 90 | Family | 0.03 | 0.42 |
| | | | | Family:Genus | 0.08 | 0.74 |
| | | | | Family:Genus:Species | 0.05 | 0.32 |
| | | | | Diet | 0.06 | 0.33 |
| | | | | Habitat | 0.01 | 0.78 |
| | | | | Diet*Habitat | 0.03 | 0.679 |
| **GME** | Full | Bray-Curtis | 94 | Family | 0.03 | 0.36 |
| | | | | Family:Genus | 0.08 | 0.60 |
| | | | | Family:Genus:Species | 0.05 | 0.34 |
| | | | | Diet | 0.07 | 0.25 |
| | | | | Habitat | 0.01 | 0.76 |
| | | | | Diet*Habitat | 0.03 | 0.60 |
| **GME** | Full | Bray-Curtis | 97 | Family | 0.03 | 0.25 |
| | | | | Family:Genus | 0.08 | 0.73 |
| | | | | Family:Genus:Species | 0.05 | 0.16 |
| | | | | Diet | 0.06 | 0.60 |
| | | | | Habitat | 0.01 | 0.61 |
| | | | | Diet*Habitat | 0.02 | 0.84 |
| **GME** | Full | Bray-Curtis | 99 | Family | 0.03 | 0.28 |
| | | | | Family:Genus | 0.08 | 0.63 |
| | | | | Family:Genus:Species | 0.05 | 0.15 |
| | | | | Diet | 0.06 | 0.65 |
| | | | | Habitat | 0.01 | 0.67 |
| | | | | Diet*Habitat | 0.02 | 0.83 |
| **GME** | Filter1 | Bray-Curtis | 90 | Family | 0.04 | 0.15 |
| | | | | Family:Genus | 0.09 | 0.35 |
| | | | | Family:Genus:Species | 0.05 | 0.20 |
| | | | | Diet | 0.07 | 0.138 |
| | | | | Habitat | 0.01 | 0.59 |
| | | | | Diet*Habitat | 0.03 | 0.26 |
| **GME** | Filter1 | Bray-Curtis | 94 | Family | 0.04 | 0.11 |
| | | | | Family:Genus | 0.09 | 0.30 |
| | | | | Family:Genus:Species | 0.05 | 0.36 |
| | | | | Diet | 0.07 | 0.13 |
| | | | | Habitat | 0.01 | 0.68 |
| | | | | Diet*Habitat | 0.03 | 0.36 |
| **GME** | Filter1 | Bray-Curtis | 97 | Family | 0.04 | 0.12 |
| | | | | Family:Genus | 0.09 | 0.50 |
| | | | | Family:Genus:Species | 0.05 | 0.22 |
| | | | | Diet | 0.06 | 0.49 |

| | | | | | | |
|---|---|---|---|---|---|---|
| | | | | Habitat | 0.01 | 0.47 |
| | | | | Diet*Habitat | 0.02 | 0.70 |
| **GME** | Filter1 | Bray-Curtis | 99 | Family | 0.04 | 0.11 |
| | | | | Family:Genus | 0.09 | 0.40 |
| | | | | Family:Genus:Species | 0.05 | 0.18 |
| | | | | Diet | 0.06 | 0.51 |
| | | | | Habitat | 0.01 | 0.50 |
| | | | | Diet*Habitat | 0.03 | 0.69 |
| **GME** | Filter2 | Bray-Curtis | 90 | Family | 0.04 | 0.17 |
| | | | | Family:Genus | 0.08 | 0.55 |
| | | | | Family:Genus:Species | 0.05 | 0.18 |
| | | | | Diet | 0.07 | 0.17 |
| | | | | Habitat | 0.01 | 0.67 |
| | | | | Diet*Habitat | 0.03 | 0.35 |
| **GME** | Filter2 | Bray-Curtis | 94 | Family | 0.04 | 0.08 |
| | | | | Family:Genus | 0.09 | 0.48 |
| | | | | Family:Genus:Species | 0.05 | 0.20 |
| | | | | Diet | 0.07 | 0.21 |
| | | | | Habitat | 0.00 | 0.83 |
| | | | | Diet*Habitat | 0.03 | 0.42 |
| **GME** | Filter2 | Bray-Curtis | 97 | Family | 0.04 | 0.12 |
| | | | | Family:Genus | 0.08 | 0.51 |
| | | | | Family:Genus:Species | 0.06 | 0.14 |
| | | | | Diet | 0.06 | 0.49 |
| | | | | Habitat | 0.01 | 0.47 |
| | | | | Diet*Habitat | 0.02 | 0.70 |
| **GME** | Filter2 | Bray-Curtis | 99 | Family | 0.04 | 0.11 |
| | | | | Family:Genus | 0.09 | 0.43 |
| | | | | Family:Genus:Species | 0.05 | 0.13 |
| | | | | Diet | 0.06 | 0.49 |
| | | | | Habitat | 0.01 | 0.54 |
| | | | | Diet*Habitat | 0.02 | 0.69 |
| **GME** | Full | Jaccard | 90 | Family | 0.03 | 0.68 |
| | | | | Family:Genus | 0.08 | 0.96 |
| | | | | Family:Genus:Species | 0.04 | 0.53 |
| | | | | Diet | 0.06 | 0.83 |
| | | | | Habitat | 0.01 | 0.63 |
| | | | | Diet*Habitat | 0.03 | 0.74 |
| **GME** | Full | Jaccard | 94 | Family | 0.03 | 0.89 |
| | | | | Family:Genus | 0.08 | 1.00 |
| | | | | Family:Genus:Species | 0.05 | 0.45 |
| | | | | Diet | 0.05 | 0.89 |
| | | | | Habitat | 0.01 | 0.87 |
| | | | | Diet*Habitat | 0.03 | 0.66 |
| **GME** | Full | Jaccard | 97 | Family | 0.03 | 0.61 |

| | | | | | | |
|---|---|---|---|---|---|---|
| | | | | Family:Genus | 0.08 | 1.00 |
| | | | | Family:Genus:Species | 0.04 | 0.65 |
| | | | | Diet | 0.06 | 0.76 |
| | | | | Habitat | 0.01 | 0.92 |
| | | | | Diet*Habitat | 0.03 | 0.94 |
| **GME** | Full | Jaccard | 99 | Family | 0.03 | 0.28 |
| | | | | Family:Genus | 0.09 | 0.57 |
| | | | | Family:Genus:Species | 0.05 | 0.13 |
| | | | | Diet | 0.06 | 0.66 |
| | | | | Habitat | 0.01 | 0.55 |
| | | | | Diet*Habitat | 0.03 | 0.44 |
| **GME** | Filter1 | Jaccard | 90 | Family | 0.03 | 0.81 |
| | | | | Family:Genus | 0.09 | 0.74 |
| | | | | Family:Genus:Species | 0.05 | 0.35 |
| | | | | Diet | 0.05 | 0.89 |
| | | | | Habitat | 0.01 | 0.85 |
| | | | | Diet*Habitat | 0.03 | 0.93 |
| **GME** | Filter1 | Jaccard | 94 | Family | 0.03 | 0.95 |
| | | | | Family:Genus | 0.08 | 0.84 |
| | | | | Family:Genus:Species | 0.04 | 0.60 |
| | | | | Diet | 0.05 | 0.91 |
| | | | | Habitat | 0.01 | 0.83 |
| | | | | Diet*Habitat | 0.03 | 0.69 |
| **GME** | Filter1 | Jaccard | 97 | Family | 0.03 | 0.49 |
| | | | | Family:Genus | 0.08 | 0.92 |
| | | | | Family:Genus:Species | 0.04 | 0.69 |
| | | | | Diet | 0.06 | 0.84 |
| | | | | Habitat | 0.01 | 0.59 |
| | | | | Diet*Habitat | 0.03 | 0.93 |
| **GME** | Filter1 | Jaccard | 99 | Family | 0.03 | 0.23 |
| | | | | Family:Genus | 0.09 | 0.39 |
| | | | | Family:Genus:Species | 0.05 | 0.17 |
| | | | | Diet | 0.06 | 0.48 |
| | | | | Habitat | 0.01 | 0.40 |
| | | | | Diet*Habitat | 0.03 | 0.45 |
| **GME** | Filter2 | Jaccard | 90 | Family | 0.03 | 0.87 |
| | | | | Family:Genus | 0.09 | 0.75 |
| | | | | Family:Genus:Species | 0.04 | 0.57 |
| | | | | Diet | 0.06 | 0.78 |
| | | | | Habitat | 0.01 | 0.84 |
| | | | | Diet*Habitat | 0.03 | 0.88 |
| **GME** | Filter2 | Jaccard | 94 | Family | 0.03 | 0.89 |
| | | | | Family:Genus | 0.08 | 0.99 |
| | | | | Family:Genus:Species | 0.05 | 0.22 |
| | | | | Diet | 0.05 | 0.97 |

| | | | | | | |
|---|---|---|---|---|---|---|
| | | | | Habitat | 0.02 | 0.37 |
| | | | | Diet*Habitat | 0.03 | 0.64 |
| **GME** | Filter2 | Jaccard | 97 | Family | 0.03 | 0.48 |
| | | | | Family:Genus | 0.08 | 0.90 |
| | | | | Family:Genus:Species | 0.04 | 0.57 |
| | | | | Diet | 0.06 | 0.77 |
| | | | | Habitat | 0.01 | 0.81 |
| | | | | Diet*Habitat | 0.03 | 0.94 |
| **GME** | Filter2 | Jaccard | 99 | Family | 0.03 | 0.19 |
| | | | | Family:Genus | 0.09 | 0.50 |
| | | | | Family:Genus:Species | 0.05 | 0.10 |
| | | | | Diet | 0.06 | 0.47 |
| | | | | Habitat | 0.01 | 0.56 |
| | | | | Diet*Habitat | 0.03 | 0.61 |
| **GME** | Full | Unweighted | 90 | Family | 0.03 | 0.76 |
| | | | | Family:Genus | 0.08 | 0.95 |
| | | | | Family:Genus:Species | 0.04 | 0.43 |
| | | | | Diet | 0.05 | 0.97 |
| | | | | Habitat | 0.02 | 0.36 |
| | | | | Diet*Habitat | 0.03 | 0.87 |
| **GME** | Full | Unweighted | 94 | Family | 0.03 | 0.76 |
| | | | | Family:Genus | 0.08 | 0.93 |
| | | | | Family:Genus:Species | 0.04 | 0.44 |
| | | | | Diet | 0.05 | 0.97 |
| | | | | Habitat | 0.02 | 0.36 |
| | | | | Diet*Habitat | 0.03 | 0.87 |
| **GME** | Full | Unweighted | 97 | Family | 0.03 | 0.57 |
| | | | | Family:Genus | 0.08 | 0.97 |
| | | | | Family:Genus:Species | 0.05 | 0.22 |
| | | | | Diet | 0.06 | 0.75 |
| | | | | Habitat | 0.01 | 0.82 |
| | | | | Diet*Habitat | 0.03 | 0.76 |
| **GME** | Full | Unweighted | 99 | Family | 0.03 | 0.36 |
| | | | | Family:Genus | 0.08 | 0.98 |
| | | | | Family:Genus:Species | 0.04 | 0.49 |
| | | | | Diet | 0.06 | 0.78 |
| | | | | Habitat | 0.01 | 0.69 |
| | | | | Diet*Habitat | 0.03 | 0.95 |
| **GME** | Filter1 | Unweighted | 90 | Family | 0.03 | 0.88 |
| | | | | Family:Genus | 0.08 | 0.79 |
| | | | | Family:Genus:Species | 0.05 | 0.33 |
| | | | | Diet | 0.05 | 0.88 |
| | | | | Habitat | 0.01 | 0.45 |
| | | | | Diet*Habitat | 0.02 | 0.94 |
| **GME** | Filter1 | Unweighted | 94 | Family | 0.03 | 0.86 |

| | | | | | | |
|---|---|---|---|---|---|---|
| | | | | Family:Genus | 0.08 | 0.80 |
| | | | | Family:Genus:Species | 0.05 | 0.33 |
| | | | | Diet | 0.05 | 0.87 |
| | | | | Habitat | 0.01 | 0.44 |
| | | | | Diet*Habitat | 0.02 | 0.95 |
| **GME** | Filter1 | Unweighted | 97 | Family | 0.03 | 0.37 |
| | | | | Family:Genus | 0.09 | 0.68 |
| | | | | Family:Genus:Species | 0.05 | 0.32 |
| | | | | Diet | 0.06 | 0.62 |
| | | | | Habitat | 0.02 | 0.27 |
| | | | | Diet*Habitat | 0.03 | 0.90 |
| **GME** | Filter1 | Unweighted | 99 | Family | 0.03 | 0.46 |
| | | | | Family:Genus | 0.08 | 0.86 |
| | | | | Family:Genus:Species | 0.05 | 0.29 |
| | | | | Diet | 0.06 | 0.59 |
| | | | | Habitat | 0.02 | 0.18 |
| | | | | Diet*Habitat | 0.02 | 0.99 |
| **GME** | Filter2 | Unweighted | 90 | Family | 0.02 | 0.93 |
| | | | | Family:Genus | 0.08 | 0.70 |
| | | | | Family:Genus:Species | 0.04 | 0.57 |
| | | | | Diet | 0.05 | 0.95 |
| | | | | Habitat | 0.01 | 0.83 |
| | | | | Diet*Habitat | 0.03 | 0.68 |
| **GME** | Filter2 | Unweighted | 94 | Family | 0.02 | 0.93 |
| | | | | Family:Genus | 0.08 | 0.71 |
| | | | | Family:Genus:Species | 0.04 | 0.63 |
| | | | | Diet | 0.05 | 0.95 |
| | | | | Habitat | 0.01 | 0.80 |
| | | | | Diet*Habitat | 0.03 | 0.65 |
| **GME** | Filter2 | Unweighted | 97 | Family | 0.03 | 0.11 |
| | | | | Family:Genus | 0.09 | 0.65 |
| | | | | Family:Genus:Species | 0.04 | 0.64 |
| | | | | Diet | 0.06 | 0.57 |
| | | | | Habitat | 0.01 | 0.84 |
| | | | | Diet*Habitat | 0.03 | 0.55 |
| **GME** | Filter2 | Unweighted | 99 | Family | 0.03 | 0.35 |
| | | | | Family:Genus | 0.08 | 0.91 |
| | | | | Family:Genus:Species | 0.05 | 0.28 |
| | | | | Diet | 0.06 | 0.53 |
| | | | | Habitat | 0.01 | 0.61 |
| | | | | Diet*Habitat | 0.03 | 0.80 |
| **GME** | Full | Weighted | 90 | Family | 0.03 | 0.29 |
| | | | | Family:Genus | 0.10 | 0.26 |
| | | | | Family:Genus:Species | 0.08 | 0.07 |
| | | | | Diet | 0.11 | 0.02 |

| | | | | | | |
|---|---|---|---|---|---|---|
| | | | | Habitat | 0.01 | 0.49 |
| | | | | Diet*Habitat | 0.02 | 0.67 |
| **GME** | Full | Weighted | 94 | Family | 0.04 | 0.22 |
| | | | | Family:Genus | 0.11 | 0.17 |
| | | | | Family:Genus:Species | 0.07 | 0.11 |
| | | | | Diet | 0.11 | 0.03 |
| | | | | Habitat | 0.01 | 0.36 |
| | | | | Diet*Habitat | 0.02 | 0.65 |
| **GME** | Full | Weighted | 97 | Family | 0.03 | 0.27 |
| | | | | Family:Genus | 0.09 | 0.28 |
| | | | | Family:Genus:Species | 0.07 | 0.10 |
| | | | | Diet | 0.10 | 0.03 |
| | | | | Habitat | 0.01 | 0.41 |
| | | | | Diet*Habitat | 0.02 | 0.60 |
| **GME** | Full | Weighted | 99 | Family | 0.02 | 0.64 |
| | | | | Family:Genus | 0.07 | 0.86 |
| | | | | Family:Genus:Species | 0.07 | 0.07 |
| | | | | Diet | 0.07 | 0.26 |
| | | | | Habitat | 0.02 | 0.32 |
| | | | | Diet*Habitat | 0.02 | 0.72 |
| **GME** | Filter1 | Weighted | 90 | Family | 0.06 | 0.05 |
| | | | | Family:Genus | 0.09 | 0.38 |
| | | | | Family:Genus:Species | 0.07 | 0.38 |
| | | | | Diet | 0.11 | 0.01 |
| | | | | Habitat | 0.01 | 0.43 |
| | | | | Diet*Habitat | 0.03 | 0.46 |
| **GME** | Filter1 | Weighted | 94 | Family | 0.06 | 0.05 |
| | | | | Family:Genus | 0.11 | 0.16 |
| | | | | Family:Genus:Species | 0.06 | 0.12 |
| | | | | Diet | 0.11 | 0.03 |
| | | | | Habitat | 0.02 | 0.24 |
| | | | | Diet*Habitat | 0.02 | 0.67 |
| **GME** | Filter1 | Weighted | 97 | Family | 0.06 | 0.04 |
| | | | | Family:Genus | 0.09 | 0.31 |
| | | | | Family:Genus:Species | 0.06 | 0.10 |
| | | | | Diet | 0.11 | 0.01 |
| | | | | Habitat | 0.02 | 0.35 |
| | | | | Diet*Habitat | 0.02 | 0.55 |
| **GME** | Filter1 | Weighted | 99 | Family | 0.03 | 0.55 |
| | | | | Family:Genus | 0.07 | 0.85 |
| | | | | Family:Genus:Species | 0.06 | 0.10 |
| | | | | Diet | 0.06 | 0.34 |
| | | | | Habitat | 0.02 | 0.20 |
| | | | | Diet*Habitat | 0.02 | 0.66 |
| **GME** | Filter2 | Weighted | 90 | Family | 0.06 | 0.04 |

| | | | | | | |
|---|---|---|---|---|---|---|
| | | | | Family:Genus | 0.08 | 0.46 |
| | | | | Family:Genus:Species | 0.07 | 0.09 |
| | | | | Diet | 0.12 | 0.02 |
| | | | | Habitat | 0.01 | 0.66 |
| | | | | Diet*Habitat | 0.03 | 0.48 |
| **GME** | Filter2 | Weighted | 94 | Family | 0.07 | 0.02 |
| | | | | Family:Genus | 0.09 | 0.35 |
| | | | | Family:Genus:Species | 0.06 | 0.14 |
| | | | | Diet | 0.11 | 0.04 |
| | | | | Habitat | 0.01 | 0.70 |
| | | | | Diet*Habitat | 0.02 | 0.66 |
| **GME** | Filter2 | Weighted | 97 | Family | 0.07 | 0.02 |
| | | | | Family:Genus | 0.08 | 0.52 |
| | | | | Family:Genus:Species | 0.06 | 0.20 |
| | | | | Diet | 0.11 | 0.02 |
| | | | | Habitat | 0.01 | 0.80 |
| | | | | Diet*Habitat | 0.03 | 0.34 |
| **GME** | Filter2 | Weighted | 99 | Family | 0.04 | 0.20 |
| | | | | Family:Genus | 0.07 | 0.84 |
| | | | | Family:Genus:Species | 0.06 | 0.19 |
| | | | | Diet | 0.07 | 0.21 |
| | | | | Habitat | 0.01 | 0.91 |
| | | | | Diet*Habitat | 0.04 | 0.17 |
| **GP** | Full | Bray-Curtis | 90 | Family | 0.29 | 0.001* |
| | | | | Family:Genus | 0.25 | 0.001* |
| | | | | Family:Genus:Species | 0.07 | 0.001* |
| | | | | Diet | 0.30 | 0.001* |
| | | | | Habitat | 0.04 | 0.001* |
| | | | | Diet*Habitat | 0.09 | 0.001* |
| **GP** | Full | Bray-Curtis | 94 | Family | 0.29 | 0.001* |
| | | | | Family:Genus | 0.25 | 0.001* |
| | | | | Family:Genus:Species | 0.07 | 0.001* |
| | | | | Diet | 0.31 | 0.001* |
| | | | | Habitat | 0.04 | 0.001* |
| | | | | Diet*Habitat | 0.09 | 0.001* |
| **GP** | Full | Bray-Curtis | 97 | Family | 0.28 | 0.001* |
| | | | | Family:Genus | 0.25 | 0.001* |
| | | | | Family:Genus:Species | 0.08 | 0.001* |
| | | | | Diet | 0.30 | 0.001* |
| | | | | Habitat | 0.04 | 0.001* |
| | | | | Diet*Habitat | 0.09 | 0.001* |
| **GP** | Full | Bray-Curtis | 99 | Family | 0.24 | 0.001* |
| | | | | Family:Genus | 0.24 | 0.001* |
| | | | | Family:Genus:Species | 0.09 | 0.001* |
| | | | | Diet | 0.27 | 0.001* |

| | | | | | | |
|---|---|---|---|---|---|---|
| | | | | Habitat | 0.05 | 0.001* |
| | | | | Diet*Habitat | 0.09 | 0.001* |
| **GP** | Full | Jaccard | 90 | Family | 0.23 | 0.001* |
| | | | | Family:Genus | 0.21 | 0.001* |
| | | | | Family:Genus:Species | 0.07 | 0.001* |
| | | | | Diet | 0.25 | 0.001* |
| | | | | Habitat | 0.03 | 0.002* |
| | | | | Diet*Habitat | 0.08 | 0.001* |
| **GP** | Full | Jaccard | 94 | Family | 0.23 | 0.001* |
| | | | | Family:Genus | 0.21 | 0.001* |
| | | | | Family:Genus:Species | 0.07 | 0.001* |
| | | | | Diet | 0.25 | 0.001* |
| | | | | Habitat | 0.03 | 0.001* |
| | | | | Diet*Habitat | 0.08 | 0.001* |
| **GP** | Full | Jaccard | 97 | Family | 0.21 | 0.001* |
| | | | | Family:Genus | 0.20 | 0.001* |
| | | | | Family:Genus:Species | 0.07 | 0.001* |
| | | | | Diet | 0.23 | 0.001* |
| | | | | Habitat | 0.04 | 0.001* |
| | | | | Diet*Habitat | 0.08 | 0.001* |
| **GP** | Full | Jaccard | 99 | Family | 0.14 | 0.001* |
| | | | | Family:Genus | 0.16 | 0.001* |
| | | | | Family:Genus:Species | 0.08 | 0.001* |
| | | | | Diet | 0.17 | 0.001* |
| | | | | Habitat | 0.03 | 0.001* |
| | | | | Diet*Habitat | 0.06 | 0.001* |
| **GP** | Full | Unweighted | 90 | Family | 0.29 | 0.001* |
| | | | | Family:Genus | 0.21 | 0.001* |
| | | | | Family:Genus:Species | 0.05 | 0.001* |
| | | | | Diet | 0.29 | 0.001* |
| | | | | Habitat | 0.03 | 0.001* |
| | | | | Diet*Habitat | 0.08 | 0.001* |
| **GP** | Full | Unweighted | 94 | Family | 0.29 | 0.001* |
| | | | | Family:Genus | 0.21 | 0.001* |
| | | | | Family:Genus:Species | 0.05 | 0.004* |
| | | | | Diet | 0.30 | 0.001* |
| | | | | Habitat | 0.03 | 0.001* |
| | | | | Diet*Habitat | 0.07 | 0.001* |
| **GP** | Full | Unweighted | 97 | Family | 0.29 | 0.001* |
| | | | | Family:Genus | 0.20 | 0.001* |
| | | | | Family:Genus:Species | 0.05 | 0.001* |
| | | | | Diet | 0.29 | 0.001* |
| | | | | Habitat | 0.03 | 0.003* |
| | | | | Diet*Habitat | 0.07 | 0.001* |
| **GP** | Full | Unweighted | 99 | Family | 0.26 | 0.001* |

| | | | | | | |
|---|---|---|---|---|---|---|
| | | | | Family:Genus | 0.20 | 0.001* |
| | | | | Family:Genus:Species | 0.06 | 0.001* |
| | | | | Diet | 0.28 | 0.001* |
| | | | | Habitat | 0.03 | 0.003* |
| | | | | Diet*Habitat | 0.07 | 0.001* |
| **GP** | Full | Weighted | 90 | Family | 0.28 | 0.001* |
| | | | | Family:Genus | 0.27 | 0.001* |
| | | | | Family:Genus:Species | 0.07 | 0.002* |
| | | | | Diet | 0.32 | 0.001* |
| | | | | Habitat | 0.03 | 0.005* |
| | | | | Diet*Habitat | 0.10 | 0.001* |
| **GP** | Full | Weighted | 94 | Family | 0.23 | 0.001* |
| | | | | Family:Genus | 0.25 | 0.001* |
| | | | | Family:Genus:Species | 0.09 | 0.001* |
| | | | | Diet | 0.27 | 0.001* |
| | | | | Habitat | 0.05 | 0.001* |
| | | | | Diet*Habitat | 0.10 | 0.001* |
| **GP** | Full | Weighted | 97 | Family | 0.25 | 0.001* |
| | | | | Family:Genus | 0.28 | 0.001* |
| | | | | Family:Genus:Species | 0.08 | 0.001* |
| | | | | Diet | 0.28 | 0.001* |
| | | | | Habitat | 0.04 | 0.005* |
| | | | | Diet*Habitat | 0.12 | 0.001* |
| **GP** | Full | Weighted | 99 | Family | 0.27 | 0.001* |
| | | | | Family:Genus | 0.27 | 0.001* |
| | | | | Family:Genus:Species | 0.06 | 0.002* |
| | | | | Diet | 0.31 | 0.001* |
| | | | | Habitat | 0.03 | 0.01* |
| | | | | Diet*Habitat | 0.10 | 0.001* |

**Table S2:** Kruskal-Wallis tests of alpha diversity for GP and GME communities. No significant associations between GME alpha diversity and ecological or taxonomic variables were detected. Significant associations were detected between GP alpha diversity, taxonomy, and diet, but not habitat.

| **Community** | **Filter** | **Metric** | **Clustering** | **Variable** | **Kruskal-Wallis** | |
|---|---|---|---|---|---|---|
| | | | | | **H-value** | **P-value** |
| **GME** | Full | Faith's PD | 90 | Family | 1.28 | 0.53 |
| | | | | Genus | 8.67 | 0.47 |
| | | | | Species | 12.39 | 0.42 |
| | | | | Diet | 5.63 | 0.34 |
| | | | | Habitat | 0.09 | 0.76 |
| **GME** | Full | Faith's PD | 94 | Family | 0.55 | 0.76 |
| | | | | Genus | 12.53 | 0.19 |
| | | | | Species | 14.28 | 0.28 |

| | | | | | | |
|---|---|---|---|---|---|---|
| | | | | Diet | 5.00 | 0.42 |
| | | | | Habitat | 0.46 | 0.50 |
| **GME** | Full | Faith’s PD | 97 | Family | 0.07 | 0.97 |
| | | | | Genus | 3.22 | 0.95 |
| | | | | Species | 6.34 | 0.90 |
| | | | | Diet | 5.46 | 0.36 |
| | | | | Habitat | 0.70 | 0.40 |
| **GME** | Full | Faith’s PD | 99 | Family | 1.40 | 0.50 |
| | | | | Genus | 10.54 | 0.31 |
| | | | | Species | 11.89 | 0.45 |
| | | | | Diet | 6.53 | 0.26 |
| | | | | Habitat | 0.13 | 0.72 |
| **GME** | Full | Obs_OTUs | 90 | Family | 0.42 | 0.81 |
| | | | | Genus | 6.65 | 0.67 |
| | | | | Species | 11.90 | 0.45 |
| | | | | Diet | 4.33 | 0.50 |
| | | | | Habitat | 0.04 | 0.84 |
| **GME** | Full | Obs_OTUs | 94 | Family | 0.01 | 1.00 |
| | | | | Genus | 6.11 | 0.73 |
| | | | | Species | 10.06 | 0.61 |
| | | | | Diet | 3.69 | 0.60 |
| | | | | Habitat | 0.00 | 0.98 |
| **GME** | Full | Obs_OTUs | 97 | Family | 0.87 | 0.65 |
| | | | | Genus | 7.54 | 0.58 |
| | | | | Species | 10.35 | 0.59 |
| | | | | Diet | 6.18 | 0.29 |
| | | | | Habitat | 0.05 | 0.83 |
| **GME** | Full | Obs_OTUs | 99 | Family | 0.46 | 0.79 |
| | | | | Genus | 5.79 | 0.76 |
| | | | | Species | 7.47 | 0.83 |
| | | | | Diet | 6.57 | 0.25 |
| | | | | Habitat | 0.28 | 0.60 |
| **GME** | Filter1 | Faith’s PD | 90 | Family | 3.57 | 0.17 |
| | | | | Genus | 12.24 | 0.20 |
| | | | | Species | 15.69 | 0.21 |
| | | | | Diet | 7.98 | 0.16 |
| | | | | Habitat | 0.10 | 0.75 |
| **GME** | Filter1 | Faith’s PD | 94 | Family | 2.62 | 0.27 |
| | | | | Genus | 14.15 | 0.12 |
| | | | | Species | 18.81 | 0.09 |
| | | | | Diet | 8.91 | 0.12 |
| | | | | Habitat | 1.71 | 0.19 |
| **GME** | Filter1 | Faith’s PD | 97 | Family | 1.08 | 0.58 |
| | | | | Genus | 6.88 | 0.65 |
| | | | | Species | 10.24 | 0.60 |

| | | | | | | |
|---|---|---|---|---|---|---|
| | | | | Diet | 5.78 | 0.33 |
| | | | | Habitat | 0.43 | 0.51 |
| **GME** | Filter1 | Faith’s PD | 99 | Family | 1.58 | 0.45 |
| | | | | Genus | 8.40 | 0.49 |
| | | | | Species | 10.79 | 0.55 |
| | | | | Diet | 5.04 | 0.41 |
| | | | | Habitat | 0.14 | 0.70 |
| **GME** | Filter1 | Obs_OTUs | 90 | Family | 0.86 | 0.65 |
| | | | | Genus | 6.23 | 0.72 |
| | | | | Species | 14.22 | 0.29 |
| | | | | Diet | 5.23 | 0.39 |
| | | | | Habitat | 0.03 | 0.86 |
| **GME** | Filter1 | Obs_OTUs | 94 | Family | 0.47 | 0.79 |
| | | | | Genus | 7.38 | 0.60 |
| | | | | Species | 13.70 | 0.32 |
| | | | | Diet | 5.69 | 0.34 |
| | | | | Habitat | 0.06 | 0.80 |
| **GME** | Filter1 | Obs_OTUs | 97 | Family | 1.22 | 0.54 |
| | | | | Genus | 6.67 | 0.67 |
| | | | | Species | 11.04 | 0.53 |
| | | | | Diet | 4.84 | 0.44 |
| | | | | Habitat | 0.34 | 0.56 |
| **GME** | Filter1 | Obs_OTUs | 99 | Family | 1.01 | 0.60 |
| | | | | Genus | 8.02 | 0.53 |
| | | | | Species | 10.34 | 0.59 |
| | | | | Diet | 6.79 | 024 |
| | | | | Habitat | 0.06 | 0.81 |
| **GME** | Filter2 | Faith’s PD | 90 | Family | 0.11 | 0.95 |
| | | | | Genus | 8.32 | 0.50 |
| | | | | Species | 11.10 | 0.52 |
| | | | | Diet | 3.32 | 0.65 |
| | | | | Habitat | 1.41 | 0.23 |
| **GME** | Filter2 | Faith’s PD | 94 | Family | 0.35 | 0.84 |
| | | | | Genus | 7.12 | 0.62 |
| | | | | Species | 13.04 | 0.37 |
| | | | | Diet | 4.25 | 0.51 |
| | | | | Habitat | 0.57 | 0.45 |
| **GME** | Filter2 | Faith’s PD | 97 | Family | 2.29 | 0.32 |
| | | | | Genus | 10.40 | 0.32 |
| | | | | Species | 12.43 | 0.41 |
| | | | | Diet | 5.14 | 0.40 |
| | | | | Habitat | 0.94 | 0.33 |
| **GME** | Filter2 | Faith’s PD | 99 | Family | 1.65 | 0.44 |
| | | | | Genus | 7.45 | 0.59 |
| | | | | Species | 10.89 | 0.54 |

| | | | | | | |
|---|---|---|---|---|---|---|
| | | | | Diet | 4.94 | 0.42 |
| | | | | Habitat | 1.17 | 0.28 |
| **GME** | Filter2 | Obs_OTUs | 90 | Family | 0.27 | 0.87 |
| | | | | Genus | 6.81 | 0.66 |
| | | | | Species | 11.60 | 0.48 |
| | | | | Diet | 2.85 | 0.72 |
| | | | | Habitat | 0.02 | 0.88 |
| **GME** | Filter2 | Obs_OTUs | 94 | Family | 0.65 | 0.72 |
| | | | | Genus | 4.89 | 0.84 |
| | | | | Species | 11.28 | 0.51 |
| | | | | Diet | 3.22 | 0.67 |
| | | | | Habitat | 0.00 | 0.95 |
| **GME** | Filter2 | Obs_OTUs | 97 | Family | 2.21 | 0.33 |
| | | | | Genus | 8.59 | 0.48 |
| | | | | Species | 11.44 | 0.49 |
| | | | | Diet | 5.62 | 0.34 |
| | | | | Habitat | 0.56 | 0.45 |
| **GME** | Filter2 | Obs_OTUs | 99 | Family | 0.34 | 0.84 |
| | | | | Genus | 8.40 | 0.49 |
| | | | | Species | 9.67 | 0.65 |
| | | | | Diet | 5.17 | 0.40 |
| | | | | Habitat | 0.09 | 0.77 |
| **GP** | Full | Faith’s PD | 90 | Family | 27.38 | 0.00* |
| | | | | Genus | 38.04 | 0.00* |
| | | | | Species | 39.98 | 0.00* |
| | | | | Diet | 25.39 | 0.00* |
| | | | | Habitat | 1.36 | 0.24 |
| **GP** | Full | Faith’s PD | 94 | Family | 25.64 | 0.00* |
| | | | | Genus | 38.03 | 0.00* |
| | | | | Species | 39.06 | 0.00* |
| | | | | Diet | 24.89 | 0.00* |
| | | | | Habitat | 1.33 | 0.25 |
| **GP** | Full | Faith’s PD | 97 | Family | 26.92 | 0.00* |
| | | | | Genus | 38.09 | 0.00* |
| | | | | Species | 38.89 | 0.00* |
| | | | | Diet | 26.08 | 0.00* |
| | | | | Habitat | 0.96 | 0.33 |
| **GP** | Full | Faith’s PD | 99 | Family | 26.67 | 0.00* |
| | | | | Genus | 38.88 | 0.00* |
| | | | | Species | 39.81 | 0.00* |
| | | | | Diet | 25.71 | 0.00* |
| | | | | Habitat | 1.40 | 0.24 |
| **GP** | Full | Obs_OTUs | 90 | Family | 27.08 | 0.00* |
| | | | | Genus | 39.46 | 0.00* |
| | | | | Species | 40.65 | 0.00* |

| | | | | | | |
|---|---|---|---|---|---|---|
| | | | | Diet | 27.10 | 0.00* |
| | | | | Habitat | 1.10 | 0.29 |
| **GP** | Full | Obs_OTUs | 94 | Family | 26.30 | 0.00* |
| | | | | Genus | 38.38 | 0.00* |
| | | | | Species | 39.60 | 0.00* |
| | | | | Diet | 26.19 | 0.00* |
| | | | | Habitat | 1.35 | 0.25 |
| **GP** | Full | Obs_OTUs | 97 | Family | 27.78 | 9.30 |
| | | | | Genus | 39.32 | 0.00* |
| | | | | Species | 40.55 | 0.00* |
| | | | | Diet | 28.03 | 0.00* |
| | | | | Habitat | 0.92 | 0.34 |
| **GP** | Full | Obs_OTUs | 99 | Family | 23.85 | 0.00* |
| | | | | Genus | 36.21 | 0.00* |
| | | | | Species | 36.97 | 0.00* |
| | | | | Diet | 23.53 | 0.00* |
| | | | | Habitat | 1.93 | 0.16 |
